# Functional characterization of N-acetyltransferase 10 (NAT10) in Leishmania mexicana

**DOI:** 10.1101/2024.09.20.614127

**Authors:** Suellen Rodrigues Maran, Ariely Barbosa Leite, Gabriela Gomes Alves, Bruno Souza Bonifácio, Carlos Eduardo Alves, Paulo Otávio Lourenco Moreira, Giovanna Marques Panessa, Heloísa Montero do Amaral Prado, Angélica Hollunder Klippel, José Renato Cussiol, Katlin Brauer Massirer, Tiago Rodrigues Ferreira, David Sacks, Clara Lúcia Barbiéri, Marcelo Santos da Silva, Rubens Lima do Monte-Neto, Nilmar Silvio Moretti

## Abstract

*Leishmania* presents a complex life cycle that involves both invertebrate and vertebrate hosts. By regulating gene expression, protein synthesis, and metabolism, the parasite can adapt to various environmental conditions. This regulation occurs mainly at the post-transcriptional level and may involve epitranscriptomic modifications of RNAs. Recent studies have shown that mRNAs in humans undergo a modification known as N4-acetylcytidine (ac4C) catalyzed by the enzyme N-acetyltransferase (NAT10), impacting mRNAs stability and translation. Here, we characterized the NAT10 homologue of *L. mexicana*, finding that the enzyme exhibits all the conserved acetyltransferase domains and although failed to functionally complement the Kre33 mutant in *Saccharomyces cerevisia*e, has *in vitro* acetyltransferase activity. We also discovered that LmexNAT10 is nuclear, and seems essential, as evidenced by unsuccessful attempts to obtain null mutant parasites. Phenotypic characterization of single-knockout parasites revealed that LmexNAT10 affects the multiplication of procyclic forms and the promastigote-amastigote differentiation. Additionally, *in vivo* infection studies using the invertebrate vector *Lutzomyia longipalpis* showed a delay in the parasite differentiation into metacyclics. Finally, we observed changes in the cell cycle progression and protein synthesis in the mutant parasites. Together, these results suggest that LmexNAT10 might be important for parasite differentiation, potentially by regulating ac4C levels.

## Introduction

Leishmaniasis is a group of diseases found in over 90 countries, manifesting in various forms depending on the *Leishmania* species involved. The main clinical forms are cutaneous, mucocutaneous, and visceral leishmaniasis – the latter being the most severe and potentially fatal if left untreated (1). Treatment is complex and prolonged, primarily relying on the use of antimony-based drugs (N-methylglucamine and sodium stibogluconate), Amphotericin B and Miltefosine (2).

*Leishmania* has a complex life cycle, alternating between invertebrate and vertebrate hosts. The invertebrate hosts are dipteran insects belonging to *Phlebotomus* and *Lutzomyia* genera, while vertebrate hosts comprise dogs, wild animals and humans. The parasite transited through three main stages: procyclic promastigote, metacyclic promastigote, and amastigote. The infection begins in the vertebrate host when it ingests amastigotes present inside macrophages of an infected mammal. In the insect digestive tract, the amastigotes are released and differentiate into procyclic promastigotes, the insect replicative form. In the next few days, sequential differentiation steps from procyclics to nectomonads and leptomonads, culminates in the rise of non-replicative metacyclics, which migrates to the anterior parts of the midgut. These metacyclic promastigotes during the insect’s feeding process are regurgitated and phagocytosed macrophages in the mammal host. Within the macrophages (inside the parasitophorous vacuole), the metacyclic promastigotes transform back into amastigotes and multiply through binary fission. This multiplication eventually causes the macrophage membrane to rupture, releasing amastigotes that infect new macrophages. The infected phagocytes can be ingested by a new vector, maintaining the infectious cycle. To survive these various environmental changes, *Leishmania* adapts by altering its biological processes, such as gene expression, protein synthesis and metabolism (3).

In *Leishmania*, most coding genes are transcribed into polycistronic mRNAs, which are later processed into monocistronic mature mRNA by trans-splicing and polyadenylation (4). During the RNA processing, a small sequence of 39 nucleotides called “spliced leader” (SL RNA) is added to 5’ end and, while a poly-A tail is added to the 3’ end of the mRNA. Due to polycistronic transcription, the regulation of gene expression is primarily regulated at the post-transcriptional level, involving mRNA degradation/stabilization and the selection of mRNA for protein synthesis (4). This suggests that chemical RNA modifications might play a significant role in gene regulation.

Chemical RNA modifications have recently emerged as a new mechanism for regulating gene expression across different organisms (5). Over 160 RNA modifications have been identified in the last years, including N6-methyladenosine (m6A), N1-methyladenosine (m1A), and N4-acetylcytidine (ac4C). Collectively referred to as epitranscriptome, these modifications influence mRNA stability, translation efficiency, and RNA-protein interactions (5,6).

The ac4C modification was first identified in yeast and mammalian tRNAs, and later in eukaryotic rRNA and bacterial tRNA (7,8). In tRNAs, it increases molecular stability and binding fidelity to the methionine-specific AUG codon. In rRNAs, this modification is important for translation and, along with other rRNA modifications, can influence the stability of interactions between ribosomal subunits(9). The presence of ac4C in the coding regions, 5’ UTRs and 3’ UTRs of mRNAs in mammals and *Saccharomyces cerevisiae* affect the stability and translation efficiency of these molecules (10). In humans, the enzyme responsible for adding ac4C to rRNA, tRNA, and mRNA is known as N-acetyltransferase 10 (NAT10). NAT10 participates in various maturation processes, including the processing of 18S rRNA and the assembly of 40S ribosomal subunit (11). Additionally, NAT10 can function as a protein N-acetyltransferase, contributing to the regulation of nuclear architecture, nucleosome structure, telomerase activity, DNA repair, cell division, spindle assembly, chromosome segregation, and stress response (12,13).

Considering the essential role of post-transcriptional regulation in *Leishmania* for controlling gene expression throughout its intricate life cycle, ac4C has the potential to play a significant role in the parasite development. In this work we characterized the function of *L. (L.) mexicana* NAT10. Specifically, we investigated the impact of this enzyme on parasite stage differentiation, cell cycle progression and protein synthesis, using single-knockout parasites.

## Results

### *In silico* analyses of LmexNAT10

To characterize the protein corresponding to NAT10 in *L. (L.) mexicana*, bioinformatics analyses were initially conducted to identify the homologous gene in this parasite. Using the amino acid sequence of human and *S. cerevisiae* NAT10, we searched the *L. (L.) mexicana* proteome and identified a homologue protein, LmxM.17.1250, with approximately 40% identity with the human NAT10 and 44% identity with the *S. cerevisia*e NAT10. This protein contains the four characteristic domains: Tmca, Helicase, GNAT, and tRNA (Figure 1A). We also identified NAT10 homologues in various *Leishmania* species, from both the *Viannia* and *Leishmania* subgenera (Figure 1A).

**Figure 1.**
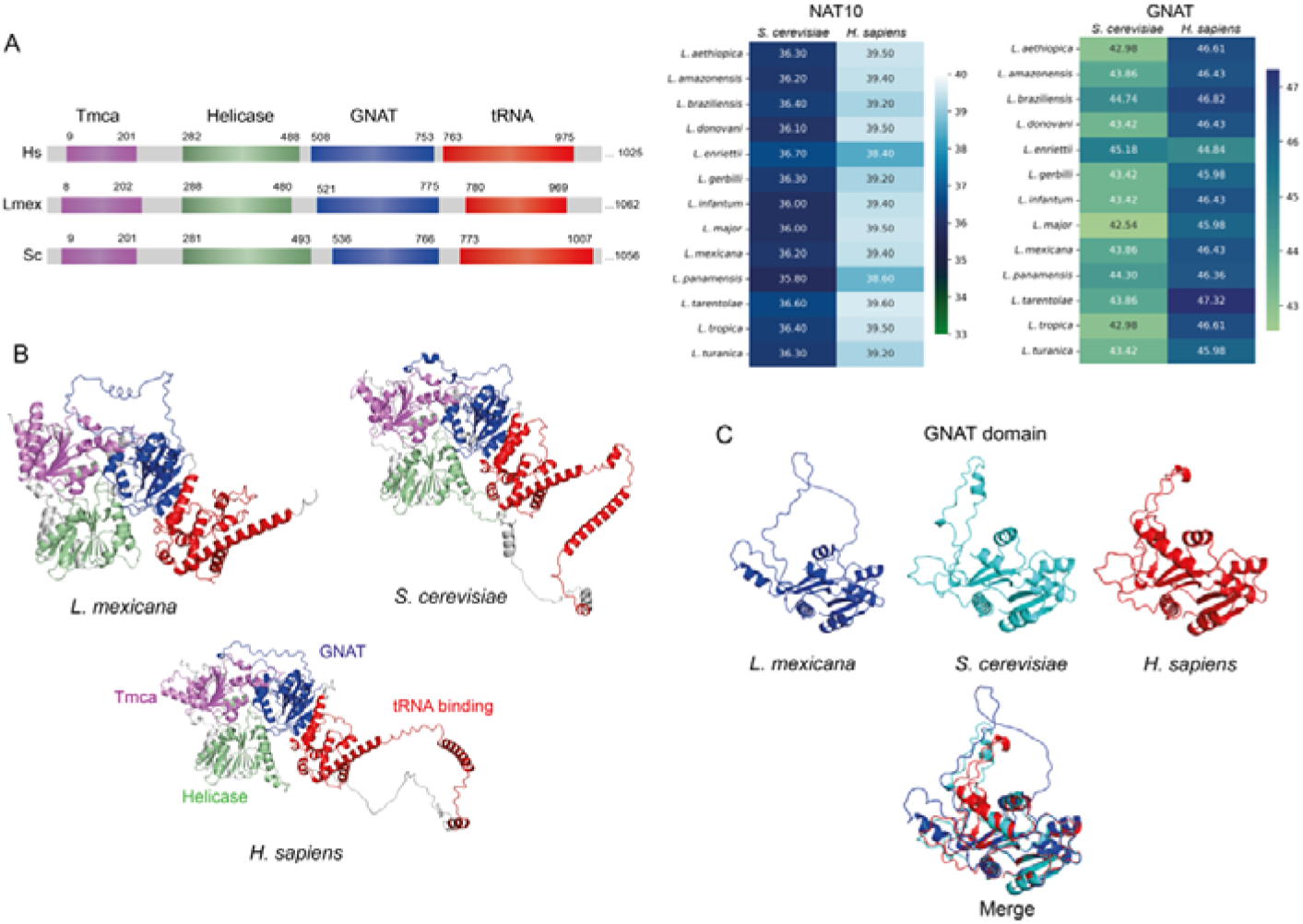
*L. (L.) mexicana* has a well-conserved NAT10 homologue. **A.** Distribution of NAT10 domains in humans, *L. mexicana*, and *S. cerevisiae*. All three species share the essential domains for NAT10 enzyme function: Tmca, Helicase, GNAT, and tRNA. The numbers above each domain indicate the start and end positions of the amino acids within each domain. Sequence identity of NAT10 between different *Leishmania* species and *S. cerevisiae* is approximately 36%, while the identity between *L. (L.) mexicana* and human NAT10 is 39.4%. The GNAT domain of *L. (L.) mexicana* shows 43.86% and 46.43% sequence identity with the corresponding domain in *S. cerevisiae* and human, respectively. **B.** Predicted 3D structures of NAT10 proteins from *L. (L.) mexicana*, *S. cerevisiae*, and human highlighting the GNAT (blue), tRNA binding (red), Tmca (purple), and Helicase (green) domains, showing that the *L. (L.) mexicana* protein has a high degree of structural conservation compared to the other species. **C.** Structural overlay of the GNAT domain shows a high degree of structural conservation among the three proteins, further illustrating the similarity in this critical functional domain.

Next, we used predicted 3D protein models of NAT10 from *L. (L.) mexicana*, *S. cerevisiae*, and human, generated with AlphaFold2, to assess the degree of structural similarity among the enzymes. We observed a high degree of structural similarity between the *L. (L.) mexicana*, human and *S. cerevisiae* proteins (Figure 1B). Specifically, when analyzing the GNAT domain - responsible for acetyltransferase activity - we also found significant structural conservation (Figure 1C).

The acetyltransferase activity of NAT10 depends on acetyl-CoA, which acts as the donor of the acetyl group that is transferred to the target molecule (protein or RNA). Therefore, we assessed the conservation of residues within the GNAT domain that are known to be involved in acetyl-CoA binding across NAT10 from *L. (L.) mexicana*, human, and *S. cerevisiae*, and we found that most of these residues conserved in the parasite protein (Figure S1).

### *S. cerevisiae* complementation assay with LmexNAT10

To begin characterizing LmexNAT10, we investigated whether the parasite gene could functionally complement a *S. cerevisiae* model. For this complementation assays, we initially attempted to generate null mutant strains of the NAT10 orthologue in yeast, called Kre33. However, we found that Kre33 is essential for *S. cerevisiae* and deletion of one copy of the gene in a diploid strain (BY4743) creating a heterozygous mutant showed no apparent phenotype when compared to the wild type strain (data not shown). As an alternative, we utilized the Kre5001 strain (14), which has a random mutation in the Kre33 gene, rendering the cell line thermosensitive at temperatures above 33°C (Figure S2A). This strain was then used for the complementation assays.

We verified the Kre5001 strain’s thermosensitive phenotype by growing it at 30 °C and 37 °C. As expected, there was a significant reduction in growth at 37 °C (Figure S2A). Next, we transformed the *kre5001* strain with the previously obtained pYES-LmexNAT10-WT and pYES-LmexNAT10-MUT constructs. The pYES vector allows the inducible expression of the protein of interest in the presence of galactose. After selecting the strains containing pYES-LmexNAT10-WT and pYES-LmexNAT10-MUT, we induced protein expression and confirmed both versions by anti-FLAG Western blot (Figure S2B). We carried out functional complementation tests using the generated strains by growing them at 30 °C and 33 °C. However, we did not observe a complete recovery of the thermosensitivity phenotype in *kre5001* at either temperature (Figure S2C). This suggests that additional factors may be required for NAT10 function in *S. cerevisiae* or that the thermosensitivity phenotype might result from secondary alterations not directly related to NAT10’s acetyltransferase function.

### LmexNAT10 exhibits *in vitro* acetyltransferase activity

To determine if LmexNAT10 exhibits acetyltransferase activity we produced heterologous protein of the wild type (LmexNAT10-WT) and its mutated form (LmexNAT10-MUT) for use in *in vitro* acetyltransferase activity assays (Figure 2A). Our results showed that LmexNAT10-WT demonstrated acetyltransferase activity, confirming its predicted function (Figure 2B).

**Figure 2.**
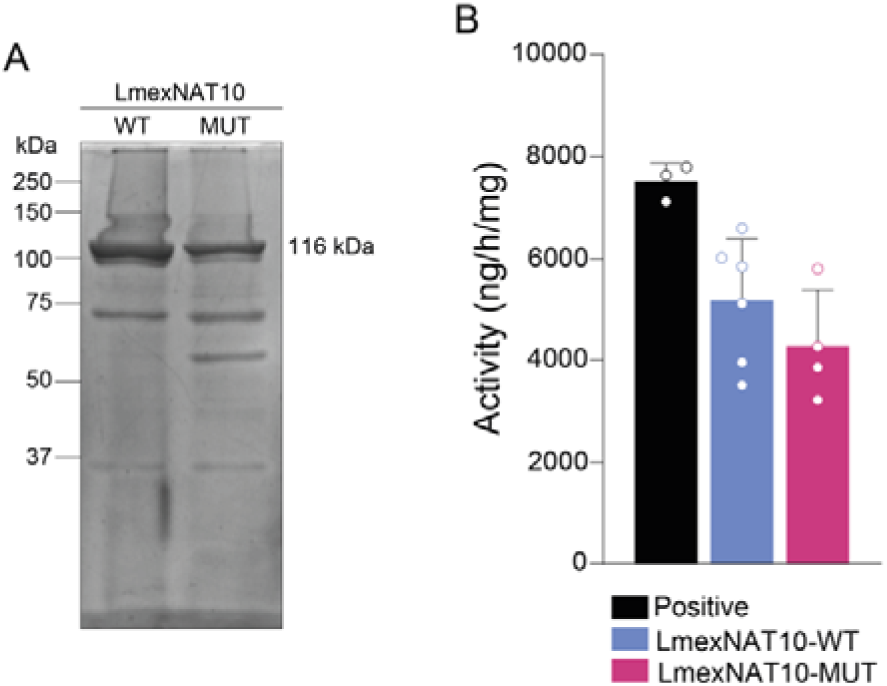
*In vitro* acetyltransferase activity of LmexNAT10. **A.** SDS-PAGE gel showing the purified LmexNAT10-WT and LmexNAT10-MUT proteins used in the acetyltransferase activity assays. The expected size of the proteins is 116 kDa. **B.** Biochemical assays using the heterologous proteins revealed that LmexNAT10-WT shows acetyltransferase activity and that the mutated protein exhibited a reduction in the activity levels. HeLA nuclear protein extracts were used as positive control. The experiments were performed in triplicate.

Additionally, we observed, a slight reduction in the activity for the LmexNAT10-MUT protein (Figure 2B).

### Generation of fluorescently tagged NAT10 and single knockout *L. (L.) mexicana* parasites

Using CRISPR-Cas9, we generated fluorescently tagged and single-allele knockout mutant parasites for the NAT10 gene in *L. (L.) mexicana*. The successful tagging of NAT10 (LmexNAT10-tag) was confirmed by flow cytometry analysis, which showed that over 90% of the LmexNAT10-tag population was fluorescence-positive compared to the parental cells (Figure 3A). To verify the correct expression of LmexNAT10 fused to mNeonGreen protein, total protein extracts of procyclic stages of both parental and LmexNAT10-tag parasites were subjected to Western blot using anti-c-Myc antibody. As expected, we detected LmexNAT10-tag only in transfected parasites, with the corrected size of 146 kDa (116 kDa + mNeonGreen + cMyc) (Figure 3B).

**Figure 3.**
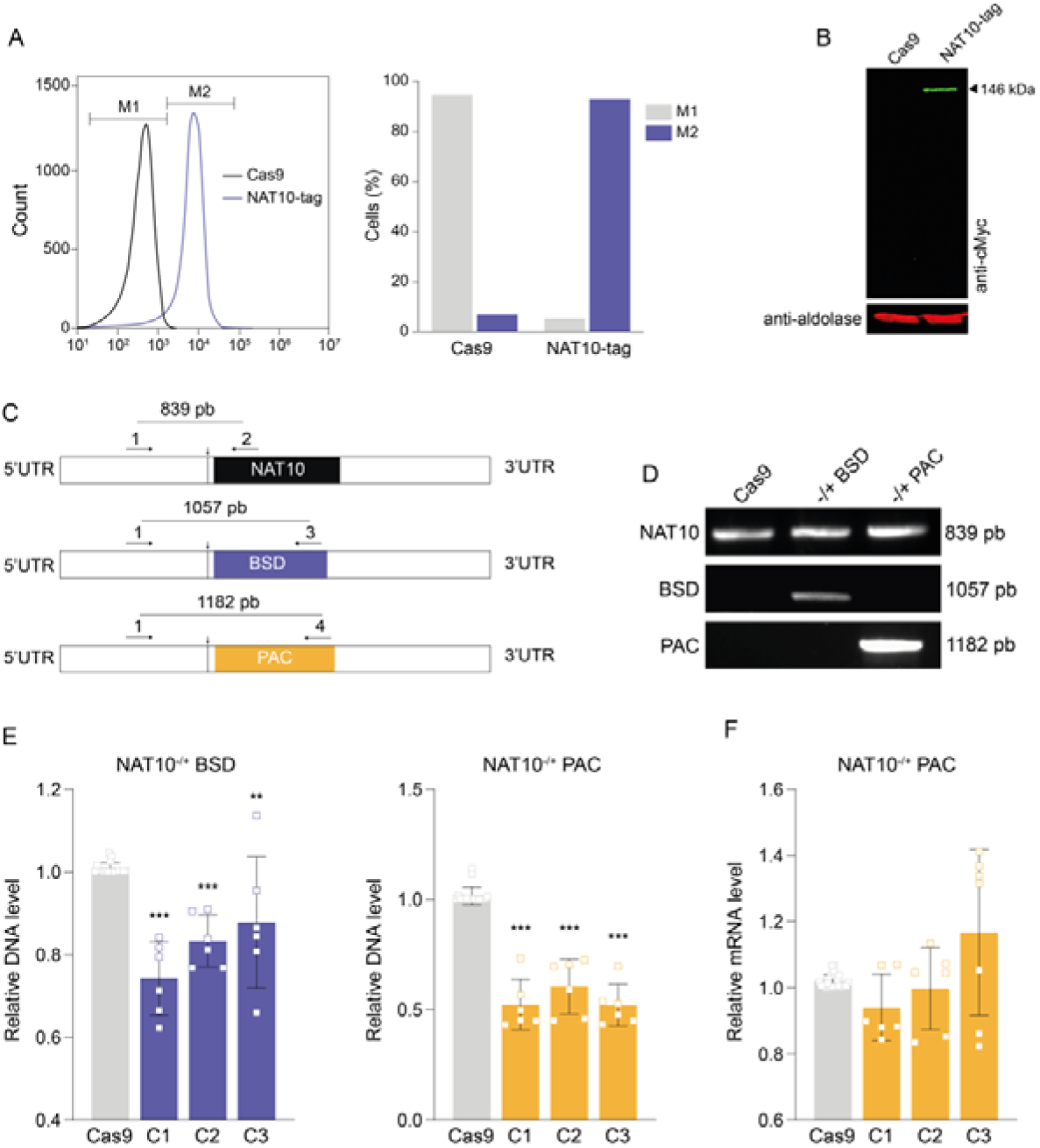
*L. (L.) mexicana* NAT10-tag and single-knockout mutant confirmation. **A**. Flow cytometry analyses of LmexNAT10-tag parasites compared to parental cells. NAT10-tag parasites exhibited higher fluorescence compared to parental cells (Cas9). **B**. Expression of LmexNAT10-tag. Extracts of total protein from LmexNAT10-tag and Cas9 cell line were used for Western blot analyses with anti-cMyc tag antibody, confirming the expression of tagged version of NAT10 in the expected size of 146 kDa. Aldolase was used as a loading control (red). **C**. Schematic representation of LmexNAT10 gene locus with the primers used for NAT10 knockout mutant confirmation. **D**. Diagnostic PCRs for LmexNAT10 single knockout. LmexNAT10 single knockout parasites were genotyped by PCR reactions using primer pairs flanking the specific locus of NAT10 (primer 1) and antibiotic-resistance genes (primer 3 or 4). **E-F**. The LmexNAT10 single knockout parasites were confirmed by qPCR and RT-qPCR reactions to measure the LmexNAT10 gene copy number, and the mRNA expression levels, respectively. Only single-knockouts (NAT10^-/+^), one allele replaced, for LmexNAT10 were obtained. All data shown are representative of three independent experiments.

After selection of LmexNAT10 knockout mutants, total genomic DNA was extracted and used to confirm the absence of NAT10 by PCR, using primers flanking the gene locus and resistance markers, as illustrated in the scheme in Figure 3C. Conventional PCR confirmed the selection of single-knockout parasites, with only one NAT10 allele replaced (Figure 3D). This result was consistently obtained after at least five rounds of transfections. Next, we selected six clones of LmexNAT10^-/+^, three from each resistance marker (BSD or PAC) and validated them by qPCR and RT-qPCR assays. First, qPCR was performed to measure NAT10 gene copy number, revealing a significant decrease for NAT10^-/+^ PAC compared to Cas9 parental cells (Figure 3E). Next, we evaluate mRNA levels in the NAT10^-/+^ PAC clone by RT-qPCR, which showed no significant difference in the mRNA (Figure 3F). These results suggested that NAT10 could be essential for procyclic promastigote stages of *L. (L.) mexicana,* as described in mammals(10).

### LmexNAT10 is a nuclear protein with developmentally regulated expression pattern

To determine the subcellular localization of LmexNAT10, we performed confocal microscopy in LmexNAT10-tagged parasites in the procyclic, metacyclic promastigotes and axenic amastigote stages. We found that NAT10 is localized in the nucleus in all parasite stages (Figure 4A and Figure S3).

**Figure 4.**
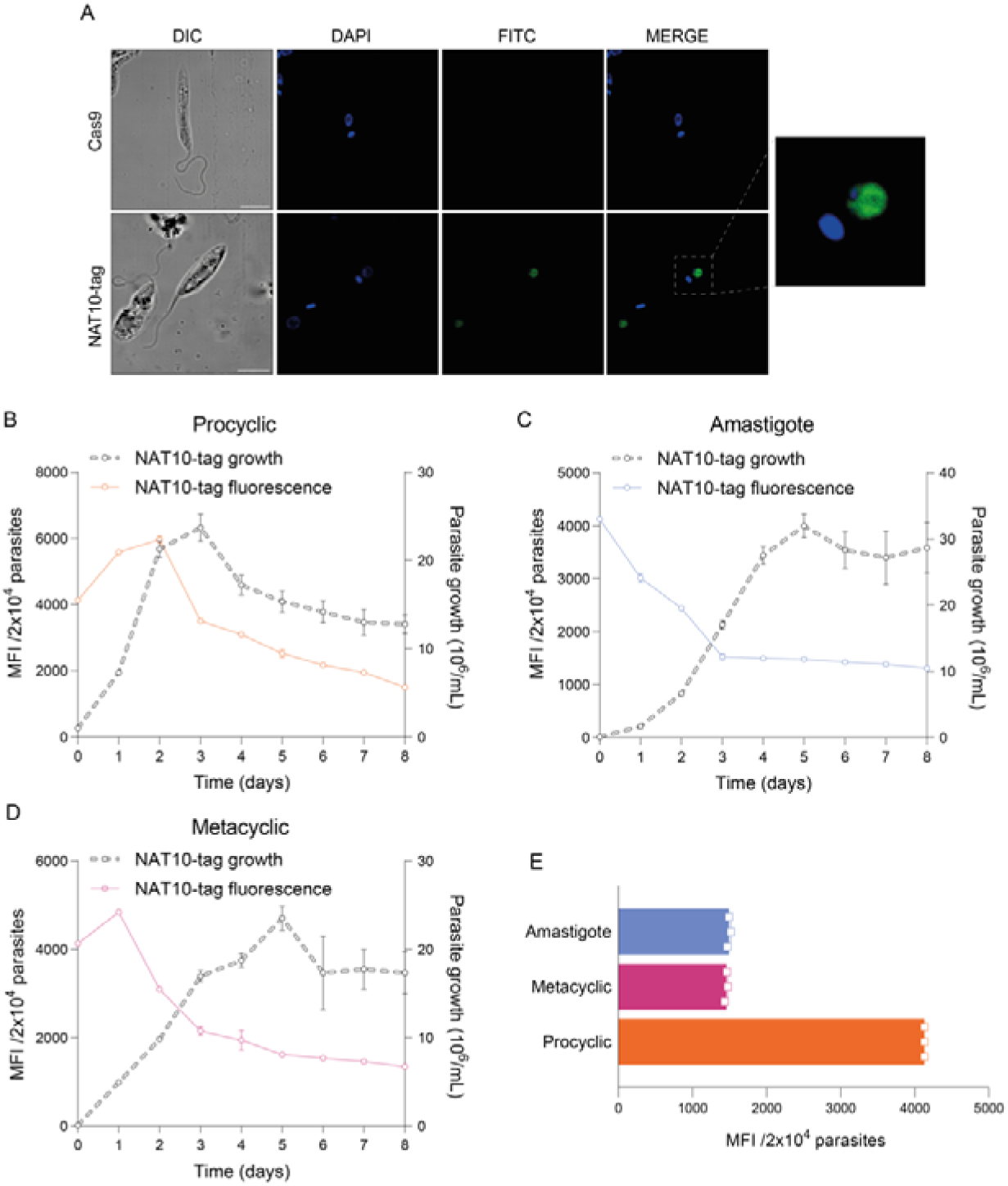
LmexNAT10 subcellular localization and expression in parasite stages. **A**. Confocal microscopy of LmexNAT10-tag procyclic stage showing its nuclear localization. Scale bar: 5 μm. **B-E**. LmexNAT10 protein indirect expression in different parasite stages. Fluorescence was measured by flow cytometry to indirectly quantify the expression of LmexNAT10 during procyclic exponential to stationary transition (B); amastigogenesis (C); metacyclogenesis (D) and in the three main parasite stages (E). Data is representative of three-independent experiments.

We also investigate whether LmexNAT10 expression varies throughout the *Leishmania* life cycle by indirect assessing protein expression during the procyclic transition from exponential to stationary phase, axenic amastigote multiplication and metacyclic differentiation. This was done by measuring fluorescence levels in parasites using flow cytometry. During the logarithmic phase of the parasite, we observed higher fluorescence levels compared to stationary phase (Figure 4B). Also, a decrease of LmexNAT10 fluorescence was noted during amastigogenesis and metacyclogenesis (Figure 4C and D), and when comparing NAT10 fluorescence across all stages, we observed high levels (4-fold time) of fluorescence in the procyclic stage (Figure 4E).

### Phenotypic analysis of LmexNAT10 single knockout parasites

After confirming the LmexNAT10^-/+^ parasites, three clones of PAC resistant were selected for phenotypic analysis experiments. To understand the role of NAT10 in the biology of *L. (L.) mexicana*, we evaluated the multiplication of procyclic, axenic amastigote differentiation, metacyclic differentiation and infection index using bone marrow-derived macrophages (BMDMs).

Initially, we monitored the *in vitro* growth of NAT10^-/+^ PAC procyclic parasites, and we observed that LmexNAT10^-/+^ PAC clones showed a slower multiplication rate compared to the counterpart parental group (Cas9), indicating that the decrease in LmexNAT10 impacts procyclic replication (Figure 5A). Next, we assessed metacyclogenesis and we found no differences in the differentiation of procyclic to metacyclic stages (Figure 5B). Conversely, the axenic and intracellular amastigote forms of LmexNAT10^-/+^ showed a significant increase in proliferation and infection when compared with Cas9 group (Figure 5C-D), suggesting that NAT10 might play different roles depending on parasite stage.

**Figure 5.**
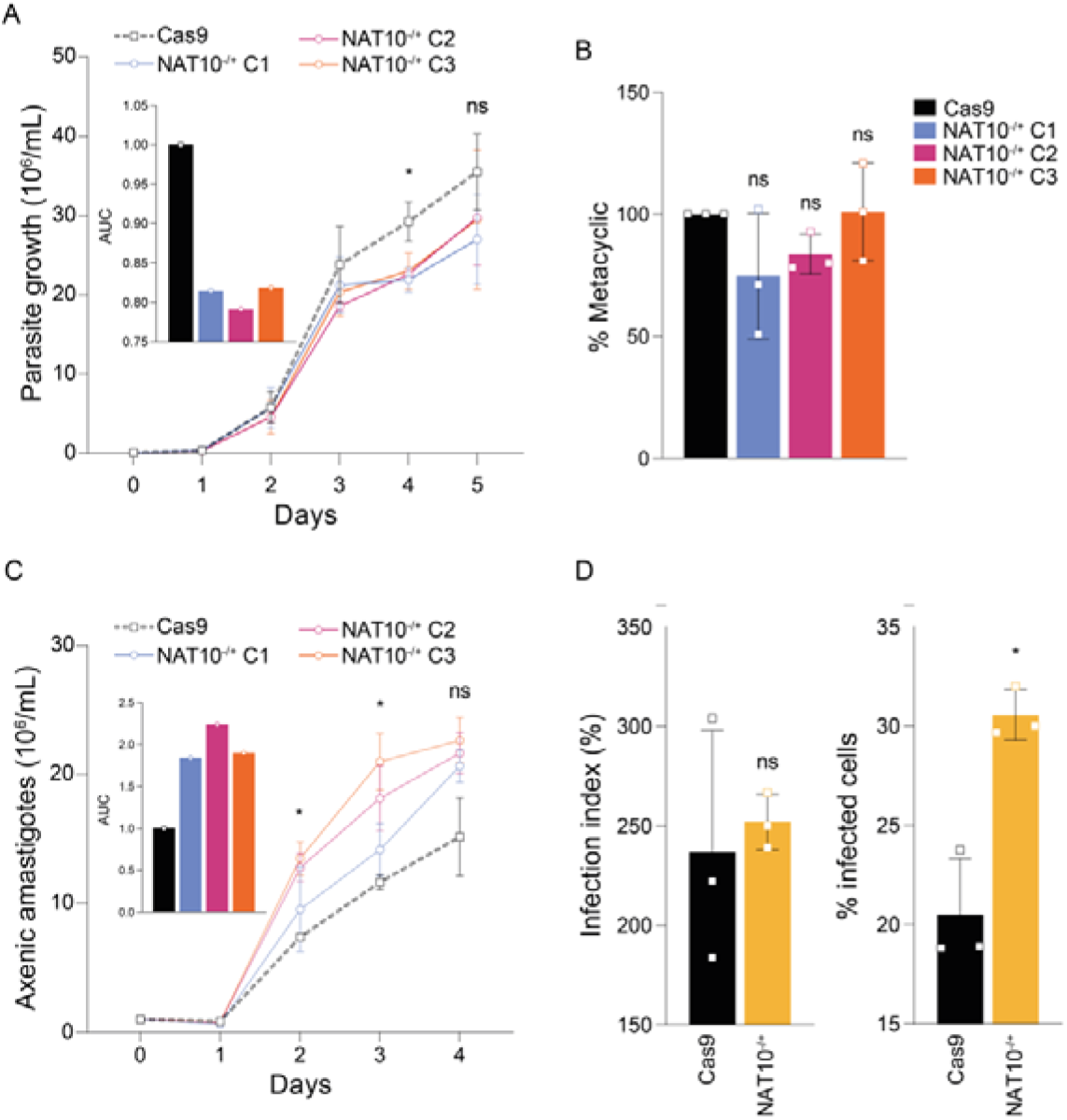
*L. mexicana* NAT10^-/+^ phenotypic analysis. **A.** LmexNAT10 affects procyclic multiplication. Growth curve and corresponding area under the curve (inside) of procyclic promastigote parental and LmexNAT10^-/+^ parasites. Statistical comparisons were made by two-way ANOVA, followed by Tukey multiple comparisons test. GP: 0.1234 (ns) and 0.0332 (*). **B.** LmexNAT10^-/+^ *in vitro* metacyclogenesis. After 7 days of metacyclogenesis in Grace’s medium, the parasites were counted before and after percoll gradient purification to determine the percentage of metacyclic promastigote. Statistical comparisons were made by one-way ANOVA followed by Dunnett’s multiple comparisons test. GP: 0.1234 (ns). **C.** Axenic amastigote differentiation and multiplication of LmexNAT10^-/+^ parasites (area under curve - AUC inset). Logarithmic procyclic promastigotes were submitted to axenic amastigote differentiation protocol and the multiplication rate was monitored daily. Statistical comparisons were made by two-way ANOVA, followed by Tukey multiple comparisons test. GP: 0.1234 (ns) and 0.0332 (*). **D.** NAT10^-/+^ *in vitro* infection with BMDMs. BMDMs isolated from BALB/c mice (1×10^5^ cells/well) were infected for 2 h with axenic amastigotes of Cas9 and LmexNAT10^-/+^ parasites. After 48 h of infection the cover glasses were fixed with methanol and stained with Giemsa. The number of amastigotes was obtained by *ImageJ software* (200 macrophages/sample). Statistical comparisons were made by one-way ANOVA with Tukey multiple comparisons test. GP:0.1234 (ns) and 0.0332 (*). All experiments were repeated three times with similar results. Three clones of NAT10 single knockout were used.

We also evaluated the metacyclogenesis *in vivo* of LmexNAT10^-/+^ compared to Cas9 using *L. longipalpis* as a model. On day 1 post-infection (p.i.) similar numbers of parasites in the sand flies infected by either cell lines were observed. At day 4 p.i. after sand fly infections by NAT10^-/+^ cells group revealed a lower frequency of metacyclics, but by day 8 p.i. the number of differentiated forms was similar between the two groups (Figure 6 and Figure S4), suggesting that NAT10 might be involved in the pre-metacyclic stage differentiation.

**Figure 6.**
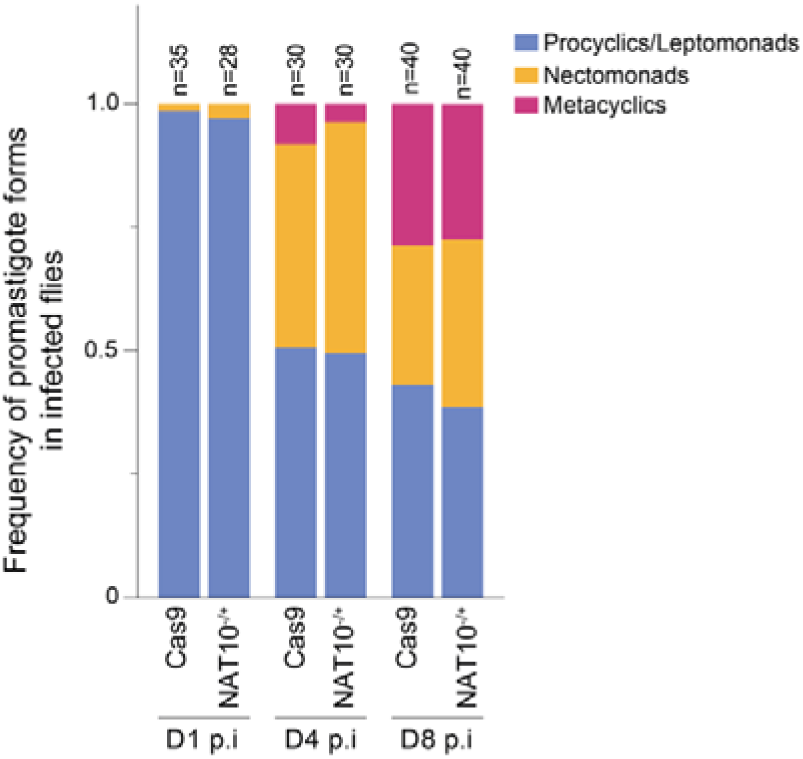
Metacyclogenesis of *L. (L.) mexicana* NAT10^-/+^ in *L. longipalpis*. *L. longipalpis* females were infected with stationary procyclic parasites of Cas9 and LmexNAT10^-/+^ and metacyclogenesis was monitored on days 1, 4 and 8 post-infections. We found that LmexNAT10^-/+^ mutants showed a delay in the differentiation process, mainly on day 4, but that it is re-established on day 8 post-infection. All infections were done at least three times with ∼100 *L. longipalpis* females for each cell line.

### LmexNAT10 affects the parasite’s cell cycle progression

Based on the results showing a delay in the LmexNAT10*^-/+^*procyclic multiplication, we decided to investigate how the absence of LmexNAT10 could impact parasite cell cycle progression. Initially we used the proliferation curves of the procyclic forms (Figure 5A) to measure the doubling time for the Cas9 line (6.25 h) and NAT10^-/+^ (6.25 h), which were then used to estimate the lengths of the cell cycle phases. Duplication time values of 6.25 and 6.25 h for the Cas9 and LmexNAT10^+/-^ strains, respectively, were found. The the percentage of cells performing cytokinesis (C) was measured. Values of 5 ± 1.3% and 5.5 ± 1.9% of procyclic 2N2K, for the Cas9 and LmexNAT10^+/^, respectively were found (Figure 7A). Therefore, C was estimated to last 0.45 h (0.07 ccu) for Cas9 and 0.49 h (0.08 ccu) for NAT10^+/-^ (Figure 7E).

**Figure 7.**
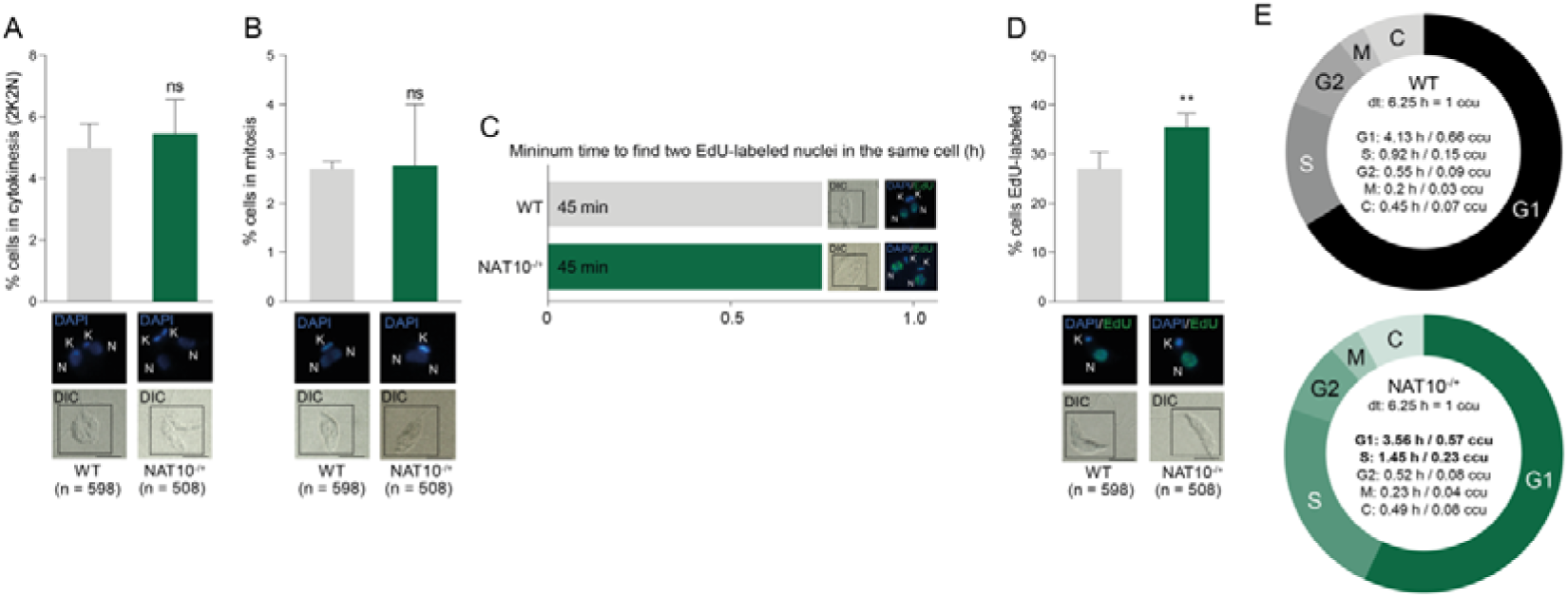
The removal of a single allele of NAT10 from *L. (L.) mexicana* affects the S phase progression. **A.** Parasites labeled with DAPI (2N2K) were used to measure the percentage of parasites in cytokinesis. **B.** Parasites with dividing nuclei, but not yet segregated, were used to estimate the proportion of parasites undergoing mitosis. **C.** To estimate the duration of the G2+M phases, EdU was added to the culture, and parasites were collected continuously every 15 min until parasites containing two EdU-labeled nuclei in the same cell (cytokinesis) were observed. This pattern was observed after 45 min for all strains. This assay was carried out in triplicate, and, in all repetitions, we found a parasite containing two EdU-labeled nuclei at the same time was found. **D.** EdU-labeled parasites were used to estimate the percentage of parasites capable of capturing this thymidine analog. The error bars in the graphs (A-D) represent the SD. NS = not significant, i.e. p > 0.05. The scale bars in the representative images (A-D) correspond to 10 µm. K = kinetoplast and N = nucleus. The values shown in each graph (A-D) represent the average of three independent assays. The number of cells analyzed (n) is shown for each strain. **E.** The measured parameters (A-D) were entered into the software CeCyD software to estimate the duration of the cell cycle phases (G1, S, G2, M and C) for each lineage analyzed. LmexNAT10-/+ presented longer S1 phase compared to Cas9 parasites. dt = doubling time; ccu = cell cycle unit.

Next, the percentage of cells in mitosis (M) was measured and values of 2.7 ± 0.26% and 2.7 ± 2% for the WT and LmexNAT10^+/-^ strains, respectively, were found (Figure 7B). Therefore, M was estimated to last 0.2 h (0.03 ccu) for Cas9 and 0.23 h (0.04 ccu) for NAT10^+/-^ (Figure 7E). In summary, there were no significant differences in the M duration between the strains. To estimate the duration of the G2+M phase, EdU was added to the culture medium, and the cells were collected continuously every 15 min (in the presence of EdU) until a cell containing two EdU-labeled nuclei (configuration 2N) to be observed. This pattern was first detected after 45 min for both Cas9 and LmexNAT10^+/-^ strains. In other words, these time values indicate that the cells at the end of S phase needed 45 min to proceed through the G2 and M phases (Figure 7C). Thus, subtracting the duration of M (previously estimated) from the duration of the G2+M phase, the G2 duration obtained was 0.55 h (0.09 ccu) for Cas9 and 0.52 h (0.08 ccu) for LmexNAT10^+/-^ (Figure 7E). As with mitosis, there was no significant differences in the duration of G2 between the two lineages analyzed. Cells labeled after 1 h with EdU pulse were used to estimate the proportion of parasites able to replicate DNA (27.1 ± 5.9% for Cas9 and 35.5 ± 4.7% for LmexNAT10^+/-^) (Figure 7D). Using this proportion and the estimated duration of the G2+M+C phases, it was possible to calculate the duration of the S phase, which was estimated to last 0.92 h (0.15 ccu) for Cas9 and 1.45 h (0.23 ccu) for LmexNAT10^+/-^ (Figure 7E). The S phase duration appears to be slightly longer for LmexNat10^+/-^ relative to control.

Finally, by subtracting the sum of C, M, G2 and S from the doubling time, we obtained the duration of the G1 phase, which was estimated at 4.13 h (0.66 ccu) for Cas9 and 3.56 h (0.57 ccu) for LmexNAT10^+/-^ (Figure 7E). Unlike the S phase duration, the G1 phase length was shown to be slightly shorter for LmexNAT10^+/-^ relative to control.

Taken together, these results suggest that the removal of a single allele of NAT10 from *L. mexicana* affects its progression in the S phase of the cell cycle, possibly delaying DNA replication, which results in a longer S phase. We believe that the observed alteration in the G1 phase duration is a consequence of the delay in DNA replication, since the other phases did not show significant differences (Figure 7E).

### Global translation is affected in NAT10^-/+^ parasites

Since the acetylation of mRNAs can increase the stability of mRNAs and translation efficiency in mammals (10), we decided to investigate protein synthesis in LmexNAT10^+/-^ parasites. Using total cell extracts, followed by sucrose gradient fractionation, we determined the polysomal profile of LmexNAT10^+/-^ and the parental Cas9 parasites, and verified a reduced amounts of polysomes in LmexNAT10^+/-^, a profile related to decreased translation initiation rates (Figure 8), compared to Cas9 cells.

**Figure 8.**
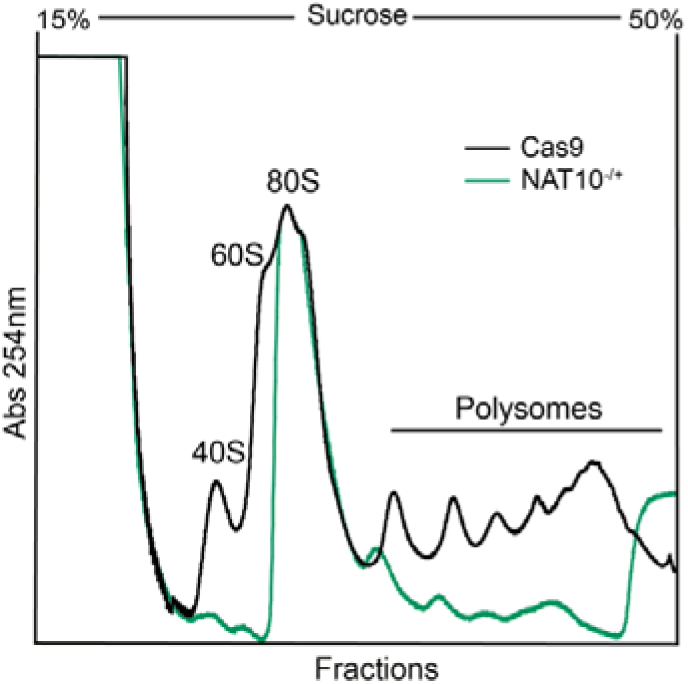
LmexNAT10-/+ procyclic parasites protein translation profile. Analysis of the polysomal profile using the Cas9 control and the LmexNAT10^+/-^ parasites in the logarithmic growth phase showed a significant reduction in the polysomes of LmexNAT10^+/-^ compared to Cas9. The position of 40S, 60S and 80S peaks and polysomes are indicated in the figure. The experiments were done at least five times and showed the similar results.

## Discussion

Regulation of *Leishmania* gene expression predominantly occurs post-transcriptionally, involving mechanisms that control mRNA stability, degradation and protein synthesis (15). This suggests that chemical RNA modifications could play an important role in this process. Here, we characterized the LmexNAT10 acetyltransferase, known to be responsible for ac4C RNA modification in various organisms and demonstrated that *L. (L.) mexicana* has a NAT10 orthologue with all domains associated with the ac4C RNA modification and acetyltransferase activity. We also found that LmexNAT10^-/+^ affects procyclic and amastigote multiplication and delayed metacyclogenesis *in vivo*, phenotypes potentially linked to disruption in cell cycle progression and protein synthesis in the parasite. Our results suggest, for the first time, that RNA modifications may be significant in *Leishmania* opening a new avenue of research to better understand the regulatory mechanism of gene expression in this parasite.

NAT10 was initially described as exhibiting histone acetyltransferase activity and a regulator of hTERT transcription, the catalytic subunit of human telomerase (16). Later studies linked NAT10 to DNA repair (13) and ac4C RNA modification in yeast and mammals (10,17). These functions are associated with the presence of four protein domains: Helicase, Tmca, GNAT and tRNA-binding.

Our bioinformatics analyses revealed that LmexNAT10 shares significant sequence identity with NAT10 from *S. cerevisiae* and human. The *Leishmania* protein contains all the essential domains and retains most of residues responsible for acetyl-CoA binding in the GNAT domain, which is linked to acetyltransferase activity. This suggests that the enzyme is fully conserved in the parasite (Figure 1).

Using the thermosensitive Kre5001 yeast cell line for complementation studies, we confirmed the expression of parasite enzyme, however, we did not observe phenotype recovery (Figure S2). The lack of recovery could be due to the requirement of *S. cerevisiae* NAT10 to interact with Tan/THUMPD1 (9,18) and LmexNAT10 may not correctly form a complex with this protein in yeast. Despite the failure of our complementation assays, we demonstrated that the heterologous LmexNAT10 possesses *in vitro* protein acetyltransferase activity (Figure 2), confirming the functional conservation of parasite enzyme.

Another conserved feature of LmexNAT10 is its nuclear localization across all parasite stages (procyclic, metacyclic promastigotes and amastigote) (Figure 4 and S3), as has been found in other organisms (16). While no changes in subcellular localization were detected, we did observe fluctuations in LmexNAT10 protein expression among the stages, with a reduction in metacyclic and amastigote compared to procyclic stage (Figure 4E). Changes in NAT10 expression has been observed under environmental stress conditions in archaea (17), suggesting that LmexNAT10 might act in parasite adaptation to different conditions found during its life cycle.

Failed attempts to generate mutant null cells demonstrated that LmexNAT10 might be essential for the parasite, consistent with findings in *S. cerevisiae* and human cells (19). The impairment of LmexNAT10^-/+^ procyclic multiplication compared to Cas9 parasites (Figure 5A), is consistent with descriptions of NAT10 in cell cycle control, as observed in mouse oocytes (20) and male germ cells (20). Furthermore, a study involving knockdown and inhibition of human NAT10 have shown that this N-acetyltransferase contributes to S phase progression, since its silencing prolongs the S phase of the cell cycle (21). It is believed that the mechanism by which NAT10 promotes cell cycle progression is via the cyclinD1/CDK2/p21 axis (21,22). These results corroborate our findings that show alterations in the cell cycle progression of LmexNAT10^-/+^, especially a delay in the S phase (Figure 7). Further studies are needed to determine the mechanism by which NAT10 is involved in the cell cycle progression of *L. mexicana* promastigote forms (Figure 7E).

On the other hand, in axenic amastigote LmexNAT10^-/+^ cells, we observed an increase in parasite multiplication. This finding is intriguing, as RNAseq data from *Trypanosoma cruzi* intracellular amastigotes showed downregulation of NAT10 during the progression of infection (23,24), suggesting that NAT10 may play different roles, depending on parasite stage.

Although we did not observe a significant effect on LmexNAT10^-/+^ metacyclogenesis, it is important to mention that for *in vivo* assays, an increased number of nectomonades was observed compared to Cas9 parasites at 1-, 4- and 8-days post-infections (Figure 6). This is particularly interesting considering given that NAT10 expression is higher in nectomonads (24,25), again indicating that this enzyme might be involved in different mechanisms depending on parasite stage.

Finally, our polysome profile data provided compelling evidence of LmexNAT10 potential impact on ac4C levels and, consequently, on protein synthesis in the parasite (Figure 8). It is well known that ac4C impacts translation efficiency in human cells (26) depending on its location within the mRNA molecule and could be functionally similar in *L. mexicana*.

In conclusion, here we showed that LmexNAT10 may play a role in regulating mechanisms involved in parasite stage differentiation, cell cycle progression and protein synthesis in *L. mexicana*. Future research should aim to confirm the presence of ac4C in *Leishmania*, determine the levels of this modification across parasite stages and explore how this might impact gene expression regulation.

## Experimental procedures

### In silico *analysis of NAT10 from* L. (L.) mexicana

For the bioinformatics analyses in this study, the amino acid sequences that code for NAT10 from of *Leishmania spp*, *S. cerevisiae*, and humans were obtained from TritrypDB (https://tritrypdb.org/tritrypdb/app) and UniProt (https://www.uniprot.org). Next, we determined the positions of the GNAT, tRNA binding, Trma, and Helicase domains for each protein using the InterPro Classification tool available on the website https://www.ebi.ac.uk/interpro/. Amino acid sequences alignment was done comparing NAT10 from *L. (L.) mexicana* with *S. cerevisiae* and humans, with the Clustal Omega tool (https://www.ebi.ac.uk/Tools/msa/clustalo/) to determine the degree of identify and the conservation of residues important for the acetyltransferase activity of *L. (L.) mexicana* NAT10. Using the sequence identity data among species, heat maps were generated with a specific pipeline created in Python (available upon request).

The predicted three-dimensional models of the proteins were obtained using the AlphaFold software, at AlphaFold Colab, available at https://colab.research.google.com/github/sokrypton/ColabFold/blob/main/AlphaFold2.ipynb. Additionally, we used the AlphaFold Protein Structure Database (https://alphafold.ebi.ac.uk). All structural analyses, such as the global structural conservation and the position of active sites for each protein were performed using Pymol 3.12 software (https://pymol.org/) and images processed with Adobe Photoshop or Adobe Illustrator.

### Constructs for *S. cerevisiae* complementation assays and protein heterologous expression

The full-length sequence corresponding to *L. (L.) mexicana* NAT10 (LmexNAT10) was obtained via PCR with specific oligonucleotides, using as a template the vector pcDNA3.1 containing the LmexNAT10, previously generated in our group. The LmexNAT10 gene was mutated at the amino acid G647 to A647 (LmexNAT10-MUT), which corresponds to an acetyl-CoA binding site described to affect NAT10 N-acetyltransferase activity.

For the amplification of LmexNAT10 (wild type - WT or mutated – MUT versions) for cloning into the pYES for *S. cerevisiae* assays, we used the oligonucleotides Forward 5’ CCGGAATTCATGGTGAAGCGCAAGGTGGA 3’ and Reverse 5’ CCGCTCGAGTCACTTGTCATCGTCGTCCT 3’ as primers, while for cloning LmexNAT10 (WT and MUT) in the pSUMO vector (27) for protein heterologous expression we used the forward oligonucleotide 5’ **ACCAGGAGCAAACGG**GAGGTATGGTGAAGCGCAAGGTGG 3’ and the reverse oligonucleotide 5’ **CAGACCGCCACCGCT**TCAACGACGCTTGGACGCCTTC 3’ - as primers. Sequences (in bold) were added to both primers to facilitate the cloning of the genes into the pSUMO expression vector by ligation-independent cloning (28,29).

For cloning in pYES backbone, the PCR reactions were carried out in a 40 μL reaction volume, with 0.5 μL of *L. (L.) mexicana* genomic DNA at a stock concentration of 4.5 μg/μL, 1 μL Taq polymerase (5 U), 3 mM MgCl_2_, 1x Taq polymerase Buffer, 0.2 μM forward and reverse primers, and 0.2 M betaine in 35 cycles with an annealing temperature of 60°C. For cloning in pSUMO backbone, the PCR reactions were carried out in a 25 μL reaction volume, with 2.5 μL of *L. mexicana* genomic DNA at a stock concentration of 2.5 μg/μL, 0.25 μL Phusion polymerase, 1x Phusion polymerase Buffer, 1 μM forward and reverse primers, in 30 cycles with touch-down PCR with annealing temperatures of 68 °C, 60 °C, 55 °C and 52 °C. The PCR products were resolved through 0.5% agarose gel electrophoresis at 120V for 45 min and used for the next cloning steps. The constructions were confirmed by endonuclease reactions and SAGER sequencing.

### S. cerevisiae complementation assays

The *S. cerevisiae* assays were performed using a conditional mutant strain *kre5001* strain, which has a mutation at the *nat10* orthologue gene, Kre33. This mutation leads to thermosensitive phenotype of the strain at temperatures higher than 30°C. This strain was used in functional complementation experiments. To confirm the thermosensitive phenotypic *kre5001* strain, a temperature sensitivity assay was performed. For the growth curves, the strains were cultured until the exponential growth phase and diluted to an O.D._600nm_ of 0.8. Subsequently, serial dilution assays were conducted in 96-well sterile plates, and the cultures were replicated using a replicator tool onto plates containing solid YPD medium. The plates were incubated at 30 °C and 37 °C for 2-3 days.

Additionally, we also used as control the wild type cell line BY4741. In both cell lines we transformed the plasmids pYES-LmexNAT10-WT and pYES-LmexNAT-MUT by lithium acetate protocol (30). For transfection, a single colony from each strain was inoculated into 5 mL of liquid YPD medium and incubated overnight at 30 °C with 250 RPM shaking. After growth, the cultures were centrifuged at 3000 x g for 4 min at 4 °C, and the pellet was resuspended in 1 mL of 100 mM LiAc. The solution was transferred to a 1.5 mL microtube and centrifuged at maximum speed for 15 sec; then the LiAc was removed. The transformation mix (PEG 50% [w/v], LiAc 1M, SS-DNA 2 mg/mL) was added to the cells and vortexed for 1 min. Subsequently, 320 μL of this solution was transferred to another tube, and 500 ng of the plasmid was added and vortexed again. The tubes were incubated at 42 °C for 40 min and then placed on ice for about 2 min. The solution was centrifuged for 10 sec, and the supernatant was removed. The cells were resuspended in 200 μL of sterile ultra-pure water and plated on selective minimal medium. Transfectants containing pYES-LmexNAT10 (WT and MUT) were plated on minimal medium (-URA) for 2-3 days at 30 °C for selection.

For thermosensitive assays, wild type and *kre5001* cell lines, bearing pYES-LmexNAT10-WT and pYES-LmexNAT10-MUT plasmids, were cultured in liquid media until the exponential growth phase and diluted to an O.D._600nm_ of 0.8. Subsequently, serial dilution assays were conducted in 96-well sterile plates, and the cultures were replicated using a replicator tool onto plates containing solid YPD medium in the presence of 2 % galactose (YPGal) or 2 % glucose (YPD). The plates were incubated at 30 °C and 33 °C for 2-3 days.

### LmexNAT10-WT and LmexNAT10-MUT heterologous expression

For bacteria heterologous protein expression, *E. coli* BL21(DE3)-R3-pRARE2 cells (31) harboring the plasmid pSUMO-LIC/LmexNAT10-WT or pSUMO-LIC/LmexNAT10-MUT were cultivated in Terrific Broth medium supplemented with 50_μg/mL kanamycin and 35_μg/mL chloramphenicol. The cells were grown under agitation (140 rpm) at 37_°C until O.D._600nm_ reached approximately 1.5. The cultures were cooled to 18 °C for 30 min, and after adding 0.2 mM isopropyl 1-thio-D-galactopyranoside to the medium, cell growth was resumed at 18 °C overnight. Cells were harvested by centrifugation and pellets were suspended in 2x lysis buffer (100 mM HEPES pH 8.0, 1.0 M NaCl, 20% glycerol, 20 mM imidazole, 2 mM PMSF, and 2 mM TCEP) in a proportion of 1 mL of buffer/g of cells. Suspended cells were flash-frozen in liquid nitrogen and stored at −80 °C. Frozen bacterial cell pellets were thawed and diluted in lysis buffer (50 mM HEPES pH 8.0, 500 mM NaCl, 10% glycerol, 10 mM imidazole, and 1 mM TCEP) in a proportion of 3 mL of buffer/g of cells and then supplemented with 0.5% CHAPS, 50 mM L-arginine and 50 mM L-glutamic acid.

Cells were lysed by sonication on ice for 4 min (5 sec on, 10 sec off, amplitude = 35%) using a Sonics Vibra Cell VCX750 ultrasonic cell disrupter. After this, 0.075% polyethyleneimine was added to the cell lysate before centrifugation at 40000 x*g* for 45 min at 4 °C. The clarified lysate was loaded onto a 5 mL HisTrap FF Crude column (GE Healthcare), equilibrated with lysis buffer, and connected to an AKTA pure protein purification system (GE Healthcare). 75 mL of wash buffer (50 mM HEPES pH 8.0, 500 mM NaCl, 10% glycerol, 30 mM imidazole, and 1 mM TCEP) was run over the column to remove any unbound protein. The His_6_/SUMO-tagged protein was eluted with 25 mL of elution buffer (50 mM HEPES pH 8.0, 500 mM NaCl, 10% glycerol, 300 mM imidazole, and 1 mM TCEP).

Removal of the His_6_/SUMO tags was performed at 4°C overnight using SUMO protease while dialyzing against excess SEC buffer (20_mM HEPES, 500_mM NaCl, 5% glycerol, and 1_mM TCEP). A second IMAC step was used to remove uncleaved protein, E. *coli* contaminant proteins, and the His_6_-tagged proteases from the cleaved protein. Qualitative analysis of the digestion reaction was performed by 12% SDS-PAGE. IMAC fractions containing the protein of interest were pooled together, concentrated to 500 µL using an Amicon® Ultra-15 15mL - 30 kDa cutoff (Merk Millipore), and injected onto a Superdex 200 Increase 10/300 (GE Life Science) column, pre-equilibrated in SEC buffer and connected to an ÄKTA Pure system (GE Healthcare) set at 0.3_mL/min. The eluate was collected in 0.4 mL fractions. SEC fractions were analyzed using 12% SDS-PAGE to confirm the purification success and determine which fractions should be pooled for final concentration. After pooling the appropriate fractions, the protein was concentrated using an Amicon® Ultra-15 15mL - 30 kDa cutoff (Merk Millipore). Protein concentration was estimated by near UV absorbance at 280 nm using a NanoDrop (Thermo Fisher Scientific). After this process, the proteins were flash-frozen in liquid nitrogen and stored at −80 °C.

### Acetyltransferase activity Assay

To perform the acetyltransferase activity assay, we used the EpiQuik™ HAT Activity/Inhibition Assay Kit (EpigenTek #P-4003), which was designed to measure the activity of histone acetylases (HAT). For each assay we used 10 µg of LmexNAT10-WT or LmexNAT10-MUT purified proteins following the manufacturer’s instructions. The assay is based on the acetylation of histone derived peptide that is detected by a high affinity anti-acetylated histone antibody. The ratio or amount of the acetylated histone, which is directly proportional to HAT enzyme activity, can then be colorimetrically quantified through an ELISA-like reaction.

### Generation of LmexNAT10 mutant cell lines

To generate *nat10* knockout and fluorescent-tagged mutants, we employed the CRISPR/Cas9 method described in (32). Specific primers were designed using the LeishGEdit tool (http://www.leishgedit.net) to generate homologous recombination fragments (HR) and to obtain the fragment for sgRNA synthesis targeting the *L. (L.) mexicana* NAT10 gene. Details of the primers used in this process can be found in Table S1. To obtain HR fragments, specific plasmids for knockout or fluorescent-tagged strategies served as templates in the PCR assays. Before transfection, the PCR products were verified for size and integrity using agarose gel electrophoresis.

For transfection, 1×10^7^ *L. (L.) mexicana* promastigotes (MHOM/GT/2001/U11032) expressing the T7 RNA polymerase and *Sp*Cas9 (T7/Cas9) at the exponential phase were resuspended in transfection cytomix buffer (66.7 mM Na_2_HPO_4_, 23.3 mM NaH_2_PO_4_, 5 mM KCl, 50 mM HEPES, pH 7.3, 150 μM CaCl_2_) and PCR products (sgRNA and HR). After that, the parasite suspension was transfected using the Amaxa NucleofectorTM IIb (Lonza). After 24 h of incubation at 26 °C, specific antibiotics were added into the culture to transfectant selection. After selection, mutant single-knockout parasites were cloned using a serial dilution method in 96-well plates and validated with PCR and RT-qPCR assays. For fluorescent-tagged mutants, flow cytometry and Western blot analysis were used to verify the indirect expression of LmexNAT10.

The LmexNAT10 single-knockout parasites were initially confirmed by PCR reactions using the genomic DNA (gDNA) as template and a combination of specific primers designed for 5’ UTR regions and for an internal region of the coding sequence of each gene. Also, specific primers for internal regions of resistance marker genes were also included in the analyses (Table S2). After the initial genotyping experiments by conventional PCR reactions, we also performed qPCR assays to quantify the gene copy number and the mRNA expression level of LmexNAT10 in the knockout mutant parasites. The gene copy number was measured using gDNA and SYBR^TM^ Green intercalator (Thermo Fisher Scientific #4367659) with specific primers for LmexNAT10. Reactions were performed in a final volume of 20 μL containing 10 pmol of forward and reverse primers, 1X SYBR^TM^ Green and 1 μL of gDNA (100 ng/μL), using the following conditions of 95 °C for 5 min followed by 45 cycles of 95 °C/15 sec, 60 °C/15 sec and 72°C/30 sec with a final extension at 72 °C for 5 min using the ViiA 7 Real-Time PCR System equipment (Thermo Fischer/Applied Biosystems). The gene copy number was calculated based on the 2^-ΔΔCT^ method and the *gapdh* gene was used as endogenous control (Table S3).

For mRNA LmexNAT10 expression levels total RNA samples from parental and mutant parasites were extracted using TRIzol method (Thermo Fisher Scientific #15596026) and treated with DNAse (Invitrogen #AM2238), prior cDNA preparation. The cDNA synthesis was done with High-Capacity cDNA Reverse Transcription Kit (Thermo Fisher Scientific #4368813) according to the manufacturer’s instructions, and a primer specific to the spliced-leader (SL) sequence was used to convert only mature mRNAs. We used the same qPCR conditions as above.

### Western blotting and flow cytometry analyses

For flow cytometry analyses 1×10^6^ cells/mL of tagged cell lines were collected and resuspended in PBS (137 mM NaCl, 2.7 mM KCl, 10 mM Na_2_HPO_4_ and 2 mM KH_2_PO_4_, pH 7.4). The samples were analyzed using the Accuri C6 flow cytometer (BD Biosciences). Analysis was conducted using the FL-1 filter and data were processed with BD Accuri^TM^ C6 software v1.0.264.21 (BD Biosciences). The parental cell was used as a negative control in the analyses.

For Western blot experiments of *L. (L.) mexicana* transfected parasites, 1×10^7^ cells were collected by centrifugation at 2500_g for 5 min, washed once in PBS and lysed using sample buffer 4x Laemmli Lysis Buffer (Sigma-Aldrich #38733). The samples were denatured at 95°C for 5 min and subjected to electrophoresis in 10% SDS-PAGE gel. The parental cells were used as a negative control in the analyses. After electrophoresis proteins were transferred to PVDF membranes, which were stained with 0.5% Ponceau S and blocked with TBSt (Tris-buffered saline with 0.05% Tween-20) and 3% skimmed milk. After that, the membranes were incubated for 1 h with anti-c-myc tag antibody (Merck #05-724) diluted [1:2500] in a blocking buffer and washed 3 times with TBSt. Next, the membranes were incubated for 1 h with secondary antibody IRDye® Goat Anti-Mouse 800CW (Biosciences 926-32210) diluted [1:10000] in a blocking buffer. To detect aldolase control load, the same steps were done but using a recombinant *T. brucei* anti-aldolase [1:5000] produced in our laboratory (33), and secondary antibody IRDye® Rabbit 680RD (Biosciences 926-68070) diluted [1:10,000] in blocking buffer. Before detection, the membranes were washed three times with TBSt and finally, the membranes were revealed using the Odissey XF® equipment (LI-COR) and the pixel quantification was performed using Image Studio version 5.2 (LI-COR Biosciences). The parental cells were used as a negative control in the analyses.

The Western blot experiments for *S. cerevisiae* were done similarly. Exponentially grown cells (wild type and *kre5001*) in the presence of galactose, were centrifuged and the resulting pellet was resuspended in TE buffer (Tris-EDTA) + 1mM PMSF. After another centrifugation, the pellet was weighed, and for every 20 mg of cells, 100 μL of lysis buffer (50 mM Tris-HCl pH 7.5; 0.2% Tergitol; 150 mM NaCl; 5 mM EDTA) + protease inhibitors were added. The cells were disrupted using a vortex mixer (Disruptor Genie - Scientific Industries) at 4 °C for 12 min and then centrifuged (4 °C/13000 x g/15 min). After lysis, 100 μL of the supernatant was mixed with 50 μL of 3X sample buffer (188 mM Tris-HCl pH 6.8; 3 % SDS; 0.01 % bromophenol blue) + DTT (25 mM) and boiled in a dry bath at 95 °C for 5 min. The remaining supernatant was used to quantify the protein concentration of the extract using the Bradford method. Subsequently, the samples were loaded onto an SDS-PAGE gel and subjected to electrophoresis (constant 100 V for 2 h). Finally, the samples were transferred to a PVDF membrane (Thermo Scientific) and incubated with primary antibodies anti-FLAG diluted 1:1000 and anti-PGK1 diluted 1:10000.

### Confocal microscopy

For the subcellular localization of all stages of LmexNAT10 fluorescent-tagged were collected and washed once with PBS. Subsequently, these cells were incubated on Poly-L-Lysine-treated slides (Tekdon Incorporated #258-041-120) for 10 min at room temperature to facilitate adherence. After cell adhesion, the wells were washed three times with PBS and fixed with 1% paraformaldehyde for 15 min at room temperature. After that, the cells were incubated with 10 μM DAPI (Invitrogen #D3571) for 10 min at room temperature, washed three times with PBS, and the slides were mounted using glycerol p-phenylenediamine (Sigma #106-50-3). The images were captured using TCS SP5 II Tandem Scanner microscopy (Leica) and processed using Imaris v6 software.

### Procyclic promastigote cultivation, metacyclogenesis and amastigogenesis assays

The procyclic stage of *L. (L.) mexicana* (MHOM/GT/2001/U11032) T7/Cas9 cell line was cultured in M199 medium (Thermo Life Science #31100019), pH 7.4, supplemented with 4.62 mM NaHCO_3_ (Sigma S5761), 40 mM HEPES (Fisher Bioreagents BP310-500), 0.1 mM adenine (Sigma-Aldrich #5251), 0.0001% biotin (Sigma-Aldrich #B329), 10 % heat-inactivated fetal calf serum (FBS) (Thermo Fisher Scientific #12657029), and 50 µg/mL hygromycin B (InvivoGen #ant-hg-5) at 26 °C. After transfection, mutant parasites were cultivated with specific selection antibiotics: 20 µg/mL blasticidin (InvivoGen #ant-bl-1) or 50 µg/mL puromycin (InvivoGen #ant-pr-5b), while fluorescent-tagged parasites were cultivated with 20 µg/mL blasticidin.

Procyclic growth curve assays were initiated by inoculating 1×10^5^ cells/mL in exponential phase into M199-supplemented medium, followed by incubation at 26 °C. Cell proliferation was monitored daily over a period of five days using the Muse Cell Analyzer (Merck Millipore).

Metacyclogenesis was induced by inoculating 1.5×10^6^ procyclic cells/mL into Grace’s medium (Sigma-Aldrich #G9771), adjusted to pH 5.5, and supplemented with 4.62 mM NaHCO3, 1X BME Vitamin (Sigma-Aldrich #B6891), 10% FBS and penicillin/streptomycin. The parasites were maintained at 26 °C for seven days to induce metacyclic stage development. Next, the parasites were resuspended in Grace’s medium and purified using a Percoll gradient (10-100%) (GE Healthcare #17089102) for metacyclic parasites purification. After that, purified metacyclics were washed with PBS and quantified using a Neubauer chamber. The metacyclogenesis rate was determined by comparing the initial number of parasites in the Percoll gradient with the final number of purified metacyclics.

The differentiation of procyclics into axenic amastigotes was initiated by incubating 1×10^6^ cells/mL in Grace’s medium, adjusted to pH 5.5 and supplemented with 4.62 mM NaHCO_3_, 1X vitamin BME (Sigma-Aldrich #B6891), and 10 % FBS, at 33°C with 5 % CO_2_ for four days. Axenic amastigotes were quantified daily using a Neubauer chamber. Additionally, the differentiation of axenic amastigotes back into procyclic promastigotes was performed inoculating 1×10^6^ cells/mL in M199-supplemented medium. Parasite growth and differentiation was monitored daily during four days by counting with a Neubauer chamber.

### In vitro macrophage infection assays

*In vitro* infection assays were done using bone marrow-derived macrophages (BMDMs) isolated from BALB/c mice. After euthanasia, the tibias and femurs were removed and disinfected with 70% ethanol. Then, the bone marrow cavities were washed using a 10 mL syringe and a 22 G1 needle with RPMI 1640 medium (Gibco #31800022) supplemented with 10% FBS. The cell population was resuspended in ACK (155 mM NH_4_Cl, 10 mM KHCO_3_ and 100 µM EDTA) and incubated at room temperature for 5 min. To obtain BMDMs cells, the population was seeded on glass coverslips and incubated in a differentiation medium (RPMI media supplemented with 10% inactivated horse serum, 30 % L929 cell conditioned medium, penicillin (100 U/mL) and streptomycin (100 µg/ml) at 37 °C with 5 % CO_2_.

After 5 days of differentiation, the macrophages were cultivated for 24 h in R10 medium and infected with axenic amastigotes MOI 1:1 (1 parasite per macrophage). After 2 h of infection, the wells were washed to remove extracellular parasites and after 12, 24 and 48 h the glass coverslips were fixed and stained with Giemsa. The images were captured with a Nikon eclipse TI-U microscope and the infection index was generated using ImageJ software.

### In vivo L. longipalpis *infection assays*

Procyclic forms of parental and LmexNAT10 single-knockout mutants were added to inactivated mice blood at a final concentration of 5×10^6^ parasites/mL, and then used to feed females of *L. longipalpis* through a chick membrane in an artificial feeder. Around 100 females of *L. longipalpis* were used per group and in the next day the fed females were transferred to a new container. To monitor the infection rate around 10 to 15 fed females were dissected and the midgut contents were extracted with PBS. The number of parasites in each midgut was accessed on days 1, 4, and 8 after infection by morphologically differentiating promastigotes/leptomonads, nectomonads and metacyclic forms as previously described in (25).

### Polysomal profile determination

Polysomal profiles were generated according to the protocol previously described by Holetz et al, 2007 (34). In summary, 1×10^9^ procyclic cells in logarithmic growth phase were incubated with 100 µg/mL cycloheximide for 10 min at 26 °C. The parasites were then centrifuged for 5 min at 2000 x g at 4 °C, washed with ice-cold PBS containing 100 µg/mL cycloheximide and resuspended in 450 µL of TKM buffer (10 mM Tris-HCl, pH 7.4, 300 mM KCl, 10 mM MgCl_2_ and 100 µg/mL cycloheximide) containing 10 µM E-64, 1 mM PMSF, 0.2 mg/mL heparin. Afterwards, the parasites were lysed by addition of 50 μL of 10 % NP-40, 2 M sucrose in TKM buffer by gentle mixing. smooth. The lysates were centrifuged for 5 min at 10000 x g at 4 °C, and the supernatant was transferred to the top of a previously prepared 15-50 % sucrose gradient in TKM buffer using a gradient generator (Gradient Master, Biocomp). The samples were ultracentrifuged at 39000 x g for 3 h in a Beckman SW41 rotor at 4 °C, and the fractions were collected and analyzed by continuous injection of 62 % sucrose at 0.6 mL/min using the Econo Gradient Pump (Bio-Rad) followed by absorbance analysis of the detector at 254 nm.

### Cell cycle analyses

To monitor DNA replication and estimate the duration of cell cycle phases, we use the thymidine analogue 5-ethynyl-2’-deoxyuridine (EdU) (ThermoFisher Scientific A10044). Procyclic of Cas9 and LmexNAT10^-/+^ cell lines were collected at logarithmic growth phase and incubated with 100 µM 5-ethynyl-2’-deoxyuridine (EdU) (ThermoFisher Scientific A10044) for different periods of time (ranging from 30 min to 1.25 h) at 26 °C. The cells were then collected (∼1×10^7^ total cells), centrifuged at 2500 g for 5 min, washed three times in PBS, fixed for 20 min with sterile 1% paraformaldehyde (Merck, 104005) diluted in PBS, washed twice and resuspended in 200 µL of PBS. Afterwards, 20 µL of the cell suspension was added to microscope slides pre-treated with microscope slides pre-treated with 0.1% poly-L-lysine and washed three times with 3% BSA (Sigma Aldrich 9048-46-8) diluted in PBS. To detect incorporated EdU, we used the Click-iT EdU detection solution for 45 min protected from light. This solution consists of 1 mM CuSO4 (Sigma Aldrich 7758-99-8), 10 µM Alexa fluor azide 488 (ThermoFisher Scientific A32723) and 100 mM C6H8O6 (Sigma Aldrich 50-81-7) diluted in distilled water (for details on the EdU procedure, see (35). The cells were then washed five times with PBS. Vectashield mounting medium (Vector H-1000-10) containing DAPI was used as a mounting solution to stain nucleus and kinetoplast DNA. The images were acquired using a fluorescence microscope (Nikon Eclipse 80i) with immersion oil at a 100x magnification. The images were analyzed with ImageJ software to count the number of EdU-positive cells in relation to the total cell number. The percentage of parasites in cytokinesis and mitosis was calculated for each group in relation to the total number of DAPI-positive parasites. When necessary, the images were superimposed using NIS elements software (Nikon).

To assess the duration of cell cycle, the profile of the nuclei (N) and kinetoplast (K) was observed in procyclic cells in logarithmic growth stained with DAPI and examined under a fluorescent microscope (Nikon). The 2N2K profile was used to estimate the percentage of cells undergoing cytokinesis (C). The cells containing a nucleus in division were used to estimate the percentage of cells in mitosis (M).

Using these values, we were able to estimate the duration of cell cycle phases cycle phases C and M according to the Williams Equation (36).

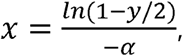

where x is the cumulative time within the cycle until the end of the stage in question, y is the cumulative % of cells including the stage in question (expressed as a fraction of a unit), and α is the specific growth rate.

To estimate the length of the G2+M phases, we added EdU to the medium containing procyclic cells in logarithmic growth and took samples every 15 min, carrying out the ‘click’ chemistry reaction, until a parasite containing two EdU-labeled nuclei (2N2K or 2N1K) was observed (this time corresponds to the length of the G2+M phases). The difference between this value and the duration of mitosis calculated above corresponds to the duration of the G2 phase.

To estimate the duration of the S phase, we measured the proportion of cells EdU-labeled cells after 1 h of EdU pulse. The duration of S phase was estimated according to the Stanners and Till equation (37):

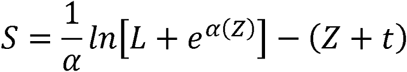

where L is the proportion of cells displaying EdU-labeled nuclei in a specific specific period, α = ln 2.T-1 (T = doubling time, expressed in hours), Z = G2 + M + cytokinesis and *t* is the duration of the EdU pulse in hours. Finally, the duration of the G1 phase was estimated by the difference between the doubling time (dt) and the sum of the other phases (S + G2 + M + C). These calculations were made using the online software CeCyD, available at https://cecyd.vital.butantan.gov.br/ (38).

## Data analysis and statistics

The results were presented as the mean ± standard deviation (SD), with the number of replicates specified for each experiment. Statistical analysis was conducted using Prism 9 (GraphPad), with significance levels indicated for each experiment.

## Acknowledgments

We are grateful to Dr. Sergio Schenkman for kindly providing access to his laboratory to carry out much of this work. We also thank Dr. Thiago Souza Onofre and Dr. Carolina Catta-Preta for their scientific support.

## Author contributions

**Suellen Rodrigues Maran:** conceptualization; data curation; formal analysis; investigation; methodology; visualization; writing – original draft. **Ariely Barbosa Leite:** conceptualization; data curation; formal analysis; investigation; visualization; writing – original draft. **Gabriela Gomes Alves:** conceptualization; data curation; formal analysis; investigation; visualization; writing – original draft. **Bruno Souza Bonifácio:** investigation; visualization. **Carlos Eduardo Gomes Alves:** investigation; visualization; software. Paulo Otávio Lourenc□**o Moreira:** investigation; visualization; formal analysis. **Giovanna Marques Panessa:** investigation; visualization. **Heloísa Montero do Amaral Prado:** investigation; visualization; conceptualization; writing – review & editing. **Angélica Hollunder Klippel:** investigation; visualization; conceptualization; writing – review & editing. **José Renato Cussiol:** conceptualization; project administration; resources; writing – review & editing. **Katlin Brauer Massirer:** conceptualization; project administration; resources; writing – review & editing. **Tiago Rodrigues Ferreira:** conceptualization; formal analysis; investigation; writing – review & editing. **David Sacks:** resources; writing – review & editing. **Clara Lúcia Barbiéri:** conceptualization; investigation; resources; writing – review & editing. **Marcelo Santos da Silva:** conceptualization; investigation; formal analysis; resources; writing – review & editing. **Rubens Lima do Monte-Neto:** conceptualization; resources; writing – review & editing. **Nilmar Silvio Moretti:** conceptualization; data curation; formal analysis; funding acquisition; investigation; methodology; resources; supervision; validation; visualization; writing – original draft; writing – review & editing.

## Ethics Statement

This research project and all the proposed procedures were submitted to and approved by the Ethics Committee on the Use of Animals and the Research Ethics Committee of the Federal University of São Paulo under registration number 9407210519/2019 and 9869091118/2019, respectively.

## Data Availability Statement

Data sharing is not applicable to this article.

---

**Figure S1.**
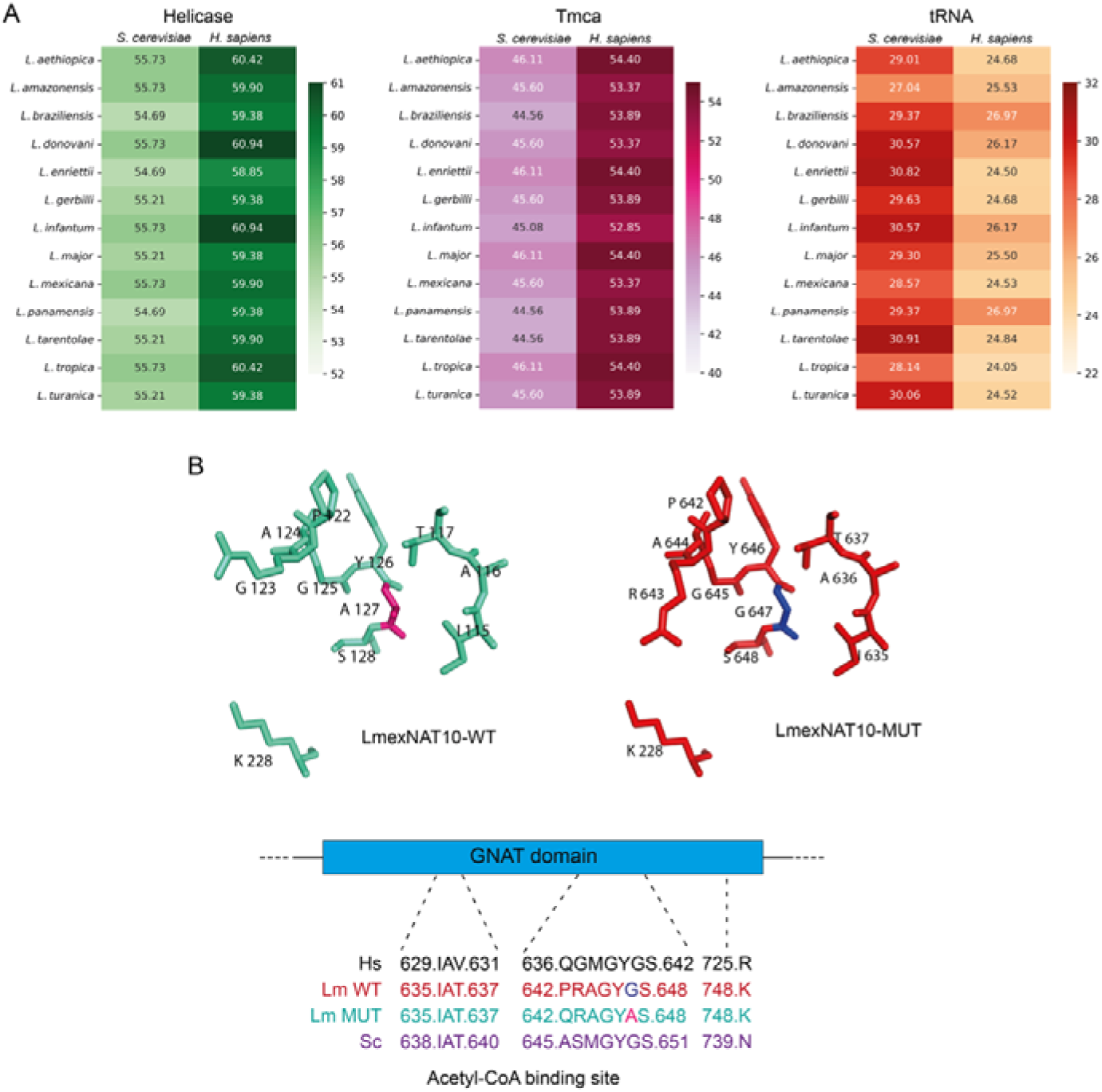
Sequence Identity of NAT10 and domains among different species of *Leishmania*, *S. cerevisiae*, and humans. **A.** The heatmaps show that the Helicase domain from *L. mexicana* and *S. cerevisiae* share 55.73% sequence identity, while *L. mexicana* and human NAT10 share ∼60%. The Tmca domain of *L. mexicana* and humans, is 53.37%. Finally, the heatmap of the tRNA domain shows that there is 28.57% identity between *L. mexicana* and *S. cerevisiae*, and 24.53% when compared to the human protein domain. **B.** The amino acid change between the wild type and mutated version of NAT10 causes a subtle alteration in the structural conformation of the GNAT domain residues. The GNAT domain residues present in the Acetyl-CoA binding site are conserved among *L. mexicana* (Lm), *S. cerevisiae* (Sc) and human (Hs).

**Figure S2.**
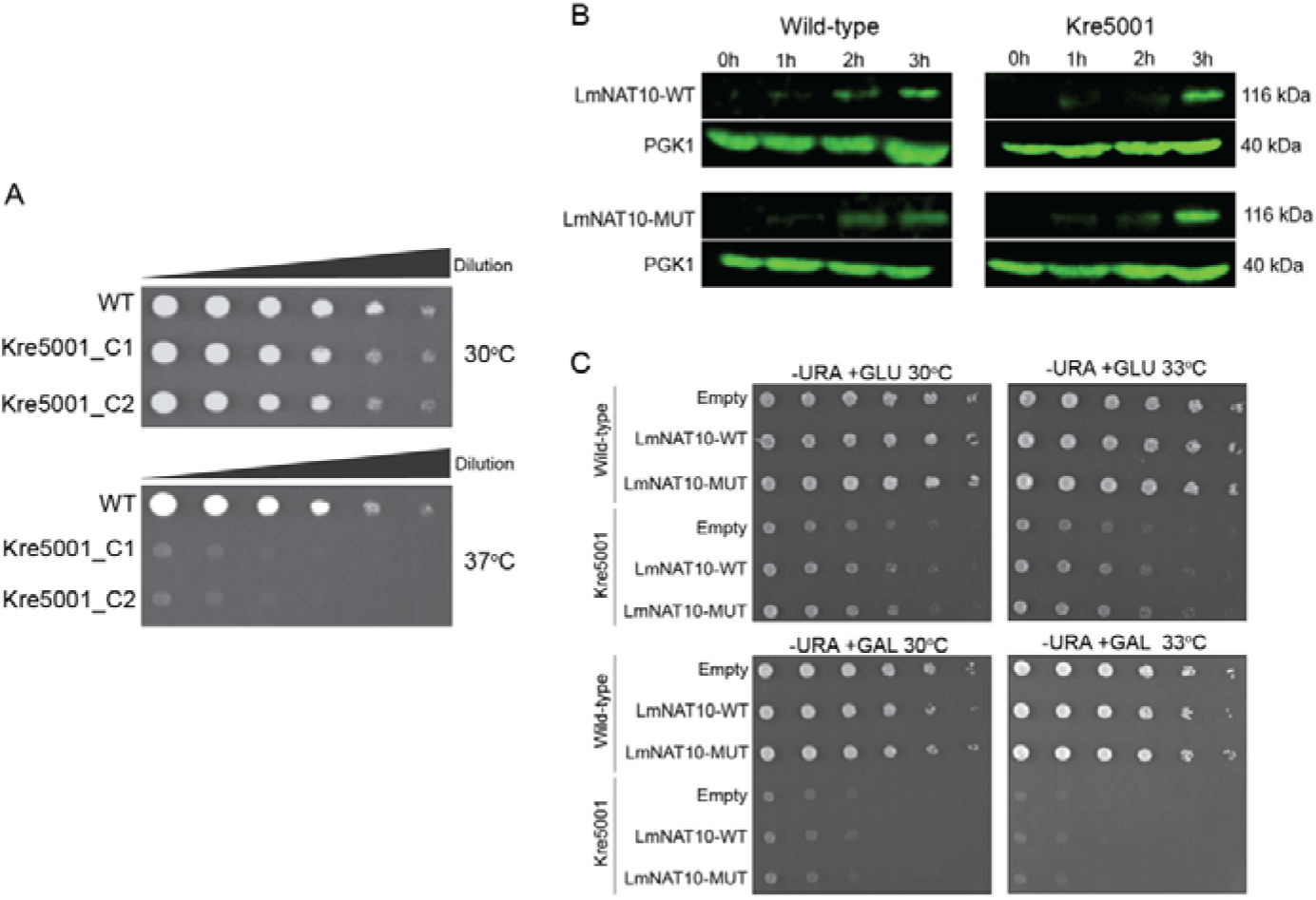
*L. mexicana* NAT10 yeast complementation assays. **A.** Wild type and Kre5001 *S. cerevisiae* strains were spotted on YPD solid medium and grown at 30°C and 37°C for 2-3 days. Growth of Kre5001 at 37°C was impaired compared to wild type. Similar growth rate was observed among the two cell lines at 30°C. **B.** Both *S. cerevisiae* strains (WT and Kre5001) were transfected with pYES-LmexNAT10-WT and pYES-LmexNAT10-MUT plasmids. Colonies were grown in liquid YPD medium containing raffinose and the plasmids were induced by galactose. Four-time points were collected (0, 1, 2, 3h) after galactose induction and Western Blotting was performed with an anti-FLAG tag antibody to confirm the expression of *L. mexicana* NAT10 protein. PGK1 protein was used as endogenous control. **C.** Thermosensitivity assays of the Kre5001 strain containing the plasmids pYES-LmNAT10-WT and pYES-LmNAT10-MUT. The Kre5001 strains pYES-LmNAT10-WT and pYES-LmNAT10-MUT were grown on solid medium in the presence or absence of galactose (GAL) at two temperatures 30°C and 33°C to assess the functional complementation of LmexNAT10. No functional complementation was observed at either temperature tested.

**Figure S3.**
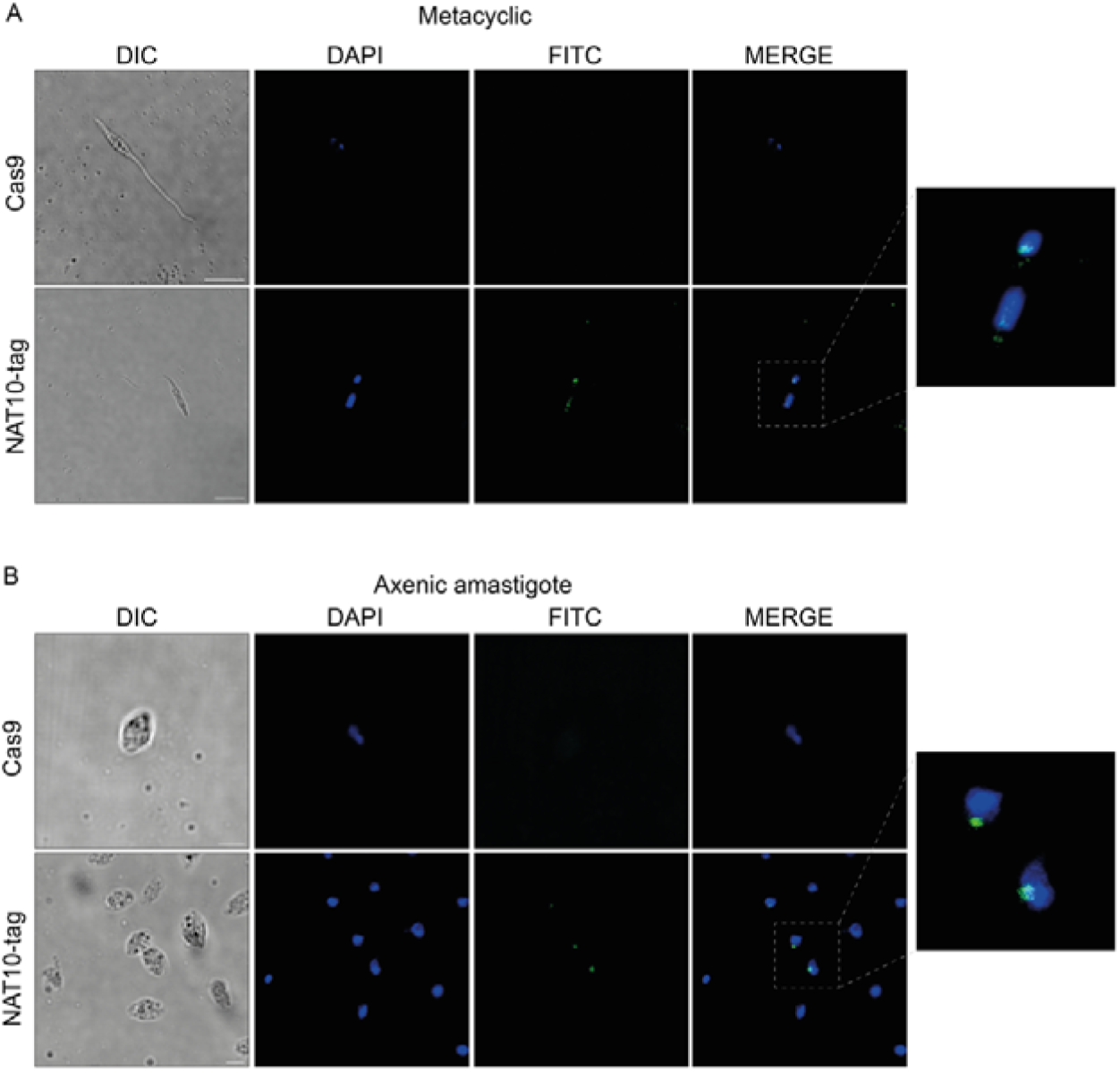
*L. mexicana* NAT10 is nuclear in metacyclic and amastigote stages. Confocal microscopy NAT10-tag cell lines of metacyclic (A) and axenic amastigote stages (B) of *L. mexicana*. Scale bars are 5 μm (metacyclic) and 2 μm (axenic amastigote).

**Figure S4.**
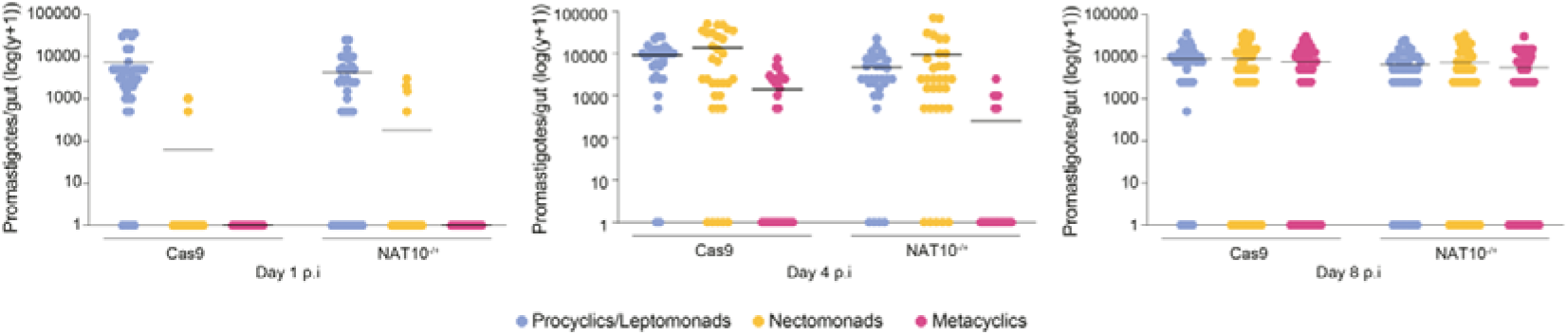
*L. mexicana* NAT10^-/+^ metacyclogenesis *in vivo*. Quantification of metacyclogenesis of Cas9 and NAT10^-/+^ at days 1, 4 and 8 post-infection of *L. longipalpis*. All infections were done at least three times with ∼100 *L. longipalpis* females for each cell line.

**Table S1.**
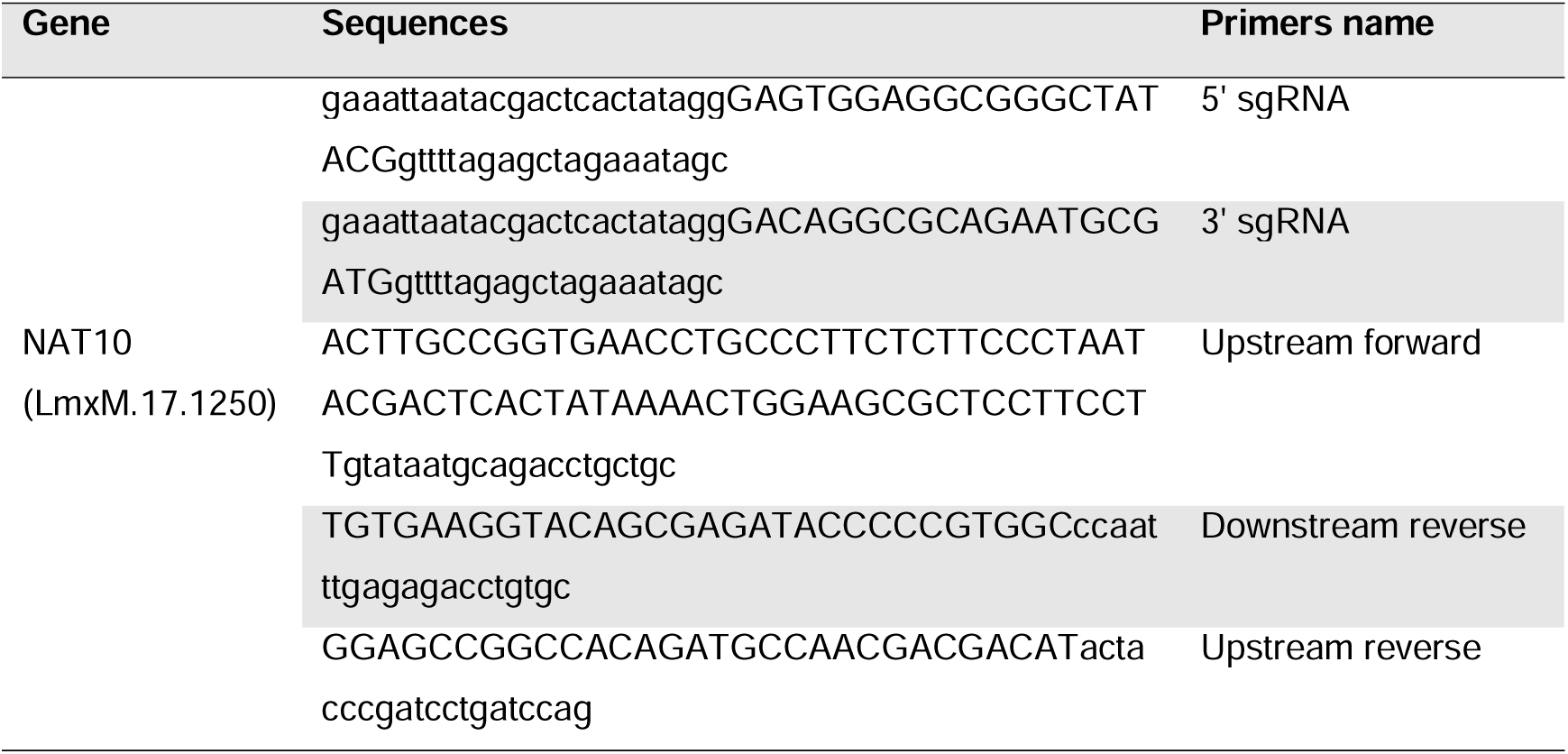
List of primers used the homologous recombination (HR) and sgRNA fragments.

**Table S2.**
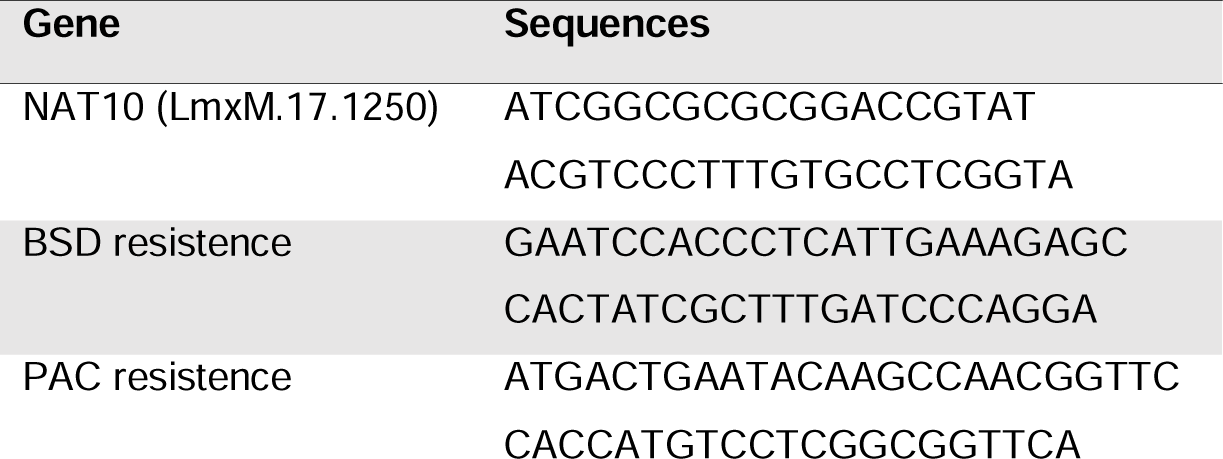
List of PCR primers for knockout parasites confirmation.

**Table S3.**
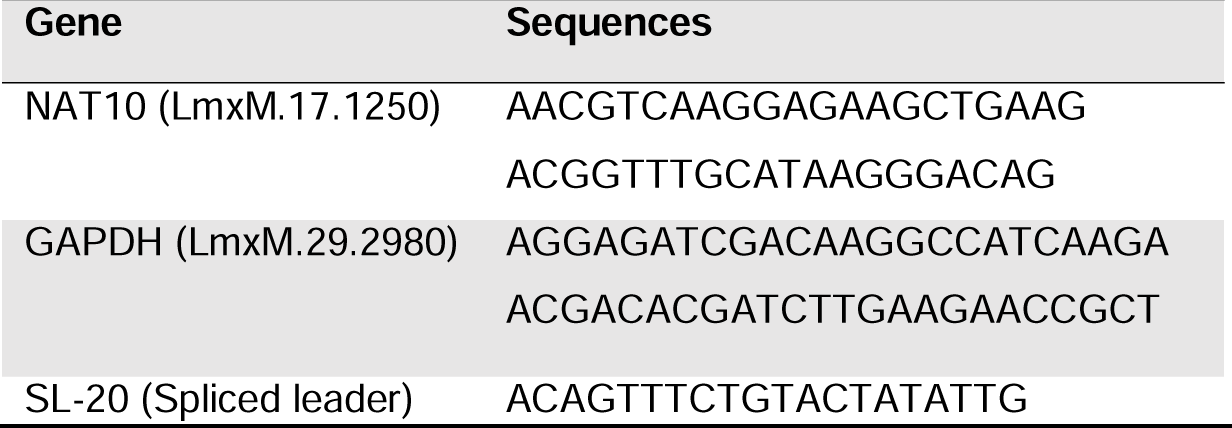
List of RT-qPCR primers for knockout parasites confirmation.

## References

1. de Vries HJC, Reedijk SH, Schallig HDFH. Cutaneous Leishmaniasis: Recent Developments in Diagnosis and Management. Vol. 16, American Journal of Clinical Dermatology. 2015.

2. Aronson N, Herwaldt BL, Libman M, Pearson R, Lopez-Velez R, Weina P, et al. Diagnosis and treatment of leishmaniasis: Clinical practice guidelines by the infectious diseases society of America (IDSA) and the American Society of tropical medicine and hygiene (ASTMH). Vol. 96, American Journal of Tropical Medicine and Hygiene. 2017.

3. Moretti NS, Schenkman S. Chromatin modifications in trypanosomes due to stress. Cell Microbiol. 2013 May;15(5):709–17.

4. De Pablos LM, Ferreira TR, Walrad PB. Developmental differentiation in Leishmania lifecycle progression: post-transcriptional control conducts the orchestra. Vol. 34, Current Opinion in Microbiology. 2016.

5. Kadumuri RV, Janga SC. Epitranscriptomic Code and Its Alterations in Human Disease. Vol. 24, Trends in Molecular Medicine. 2018.

6. Gilbert W V., Bell TA, Schaening C. Messenger RNA modifications: Form, distribution, and function. Science (1979). 2016;352(6292).

7. Bruenger E, Kowalak JA, Kuchino Y, McCloskey JA, Mizushima H, Stetter KO, et al. 5S rRNA modification in the hyperthermophilic archaea Sulfolobus solfataricus and Pyrodictium occultum. The FASEB Journal. 1993;7(1).

8. Staehelin M, Rogg H, Baguley BC, Ginsberg T, Wehrli W. Structure of a mammalian serine tRNA. Vol. 219, Nature. 1968.

9. Sharma S, Langhendries JL, Watzinger P, Kotter P, Entian KD, Lafontaine DLJ. Yeast Kre33 and human NAT10 are conserved 18S rRNA cytosine acetyltransferases that modify tRNAs assisted by the adaptor Tan1/THUMPD1. Nucleic Acids Res. 2015;43(4).

10. Arango D, Sturgill D, Alhusaini N, Dillman AA, Sweet TJ, Hanson G, et al. Acetylation of Cytidine in mRNA Promotes Translation Efficiency. Cell. 2018;175(7).

11. Sleiman S, Dragon F. Recent advances on the structure and function of RNA acetyltransferase Kre33/NAT10. Vol. 8, Cells. 2019.

12. Shen Q, Zheng X, McNutt MA, Guang L, Sun Y, Wang J, et al. NAT10, a nucleolar protein, localizes to the midbody and regulates cytokinesis and acetylation of microtubules. Exp Cell Res. 2009;315(10).

13. Liu H, Ling Y, Gong Y, Sun Y, Hou L, Zhang B. DNA damage induces N-acetyltransferase NAT10 gene expression through transcriptional activation. Mol Cell Biochem. 2007;300(1–2).

14. Costanzo M, VanderSluis B, Koch EN, Baryshnikova A, Pons C, Tan G, et al. A global genetic interaction network maps a wiring diagram of cellular function. Science (1979). 2016;353(6306).

15. Clayton C, Shapira M. Post-transcriptional regulation of gene expression in trypanosomes and leishmanias. Vol. 156, Molecular and Biochemical Parasitology. 2007.

16. Lv J, Liu H, Wang Q, Tang Z, Hou L, Zhang B. Molecular cloning of a novel human gene encoding histone acetyltransferase-like protein involved in transcriptional activation of hTERT. Biochem Biophys Res Commun. 2003;311(2).

17. Sas-Chen A, Thomas JM, Matzov D, Taoka M, Nance KD, Nir R, et al. Dynamic RNA acetylation revealed by quantitative cross-evolutionary mapping. Nature. 2020;583(7817).

18. Ma CR, Liu N, Li H, Xu H, Zhou XL. Activity reconstitution of Kre33 and Tan1 reveals a molecular ruler mechanism in eukaryotic tRNA acetylation. Nucleic Acids Res. 2024 May 22;52(9):5226–40.

19. Thalalla Gamage S, Howpay Manage SA, Chu TT, Meier JL. Cytidine Acetylation Across the Tree of Life. Acc Chem Res. 2024;57(3).

20. Chen L, Wang WJ, Liu Q, Wu YK, Wu YW, Jiang Y, et al. NAT10-mediated N4-acetylcytidine modification is required for meiosis entry and progression in male germ cells. Nucleic Acids Res. 2022;50(19).

21. Oh TI, Lee YM, Lim BO, Lim JH. Inhibition of NAT10 suppresses melanogenesis and melanoma growth by attenuating microphthalmia-associated transcription factor (MITF) expression. Int J Mol Sci. 2017;18(9).

22. Dalhat MH, Altayb HN, Khan MI, Choudhry H. Structural insights of human N-acetyltransferase 10 and identification of its potential novel inhibitors. Sci Rep. 2021;11(1).

23. Li Y, Shah-Simpson S, Okrah K, Belew AT, Choi J, Caradonna KL, et al. Transcriptome Remodeling in Trypanosoma cruzi and Human Cells during Intracellular Infection. PLoS Pathog. 2016;12(4).

24. Maran SR, de Lemos Padilha Pitta JL, dos Santos Vasconcelos CR, McDermott SM, Rezende AM, Silvio Moretti N. Epitranscriptome machinery in Trypanosomatids: New players on the table? Mol Microbiol. 2021;115(5).

25. Inbar E, Hughitt VK, Dillon LAL, Ghosh K, El-Sayed NM, Sacks DL. The transcriptome of Leishmania major developmental stages in their natural sand fly vector. mBio. 2017;8(2).

26. Arango D, Sturgill D, Yang R, Kanai T, Bauer P, Roy J, et al. Direct epitranscriptomic regulation of mammalian translation initiation through N4-acetylcytidine. Mol Cell. 2022;82(15).

27. Weeks SD, Drinker M, Loll PJ. Ligation independent cloning vectors for expression of SUMO fusions. Protein Expr Purif. 2007;53(1).

28. Tosarini TR, Ramos PZ, Profeta GS, Baroni RM, Massirer KB, Couñago RM, et al. Cloning, expression and purification of kinase domains of cacao PR-1 receptor-like kinases. Protein Expr Purif. 2018;146.

29. Savitsky P, Bray J, Cooper CDO, Marsden BD, Mahajan P, Burgess-Brown NA, et al. High-throughput production of human proteins for crystallization: The SGC experience. J Struct Biol. 2010;172(1).

30. Gietz RD, Woods RA. Yeast transformation by the LiAc/SS Carrier DNA/PEG method. Methods Mol Biol. 2006;313.

31. Gileadi O, Burgess-Brown NA, Colebrook SM, Berridge G, Savitsky P, Smee CEA, et al. High throughput production of recombinant human proteins for crystallography. Methods Mol Biol. 2008;426.

32. Beneke T, Madden R, Makin L, Valli J, Sunter J, Gluenz E. A CRISPR Cas9 high-throughput genome editing toolkit for kinetoplastids. R Soc Open Sci. 2017;4(5).

33. Leite AB, Gomes AAS, Sousa ACDCN, Fontes MRDM, Schenkman S, Moretti NS. Effect of lysine acetylation on the regulation of Trypanosoma brucei glycosomal aldolase activity. Biochemical Journal. 2020;477(9).

34. Holetz FB, Correa A, Ávila AR, Nakamura CV, Krieger MA, Goldenberg S. Evidence of P-body-like structures in Trypanosoma cruzi. Biochem Biophys Res Commun. 2007;356(4).

35. Salic A, Mitchison TJ. A chemical method for fast and sensitive detection of DNA synthesis in vivo. Proc Natl Acad Sci U S A. 2008;105(7).

36. Williams FM. A model of cell growth dynamics. J Theor Biol. 1967;15(2).

37. Stanners CP, Till JE. DNA synthesis in individual L-strain mouse cells. BBA - Biochimica et Biophysica Acta. 1960;37(3).

38. da Silva MS, Muñoz PAM, Armelin HA, Elias MC. Differences in the Detection of BrdU/EdU Incorporation Assays Alter the Calculation for G1, S, and G2 Phases of the Cell Cycle in Trypanosomatids. Journal of Eukaryotic Microbiology. 2017;64(6).

