## Supplementary files for "Functional characterization of N-acetyltransferase 10 (NAT10) in Leishmania mexicana"

**Sup**p**lementary Figures**

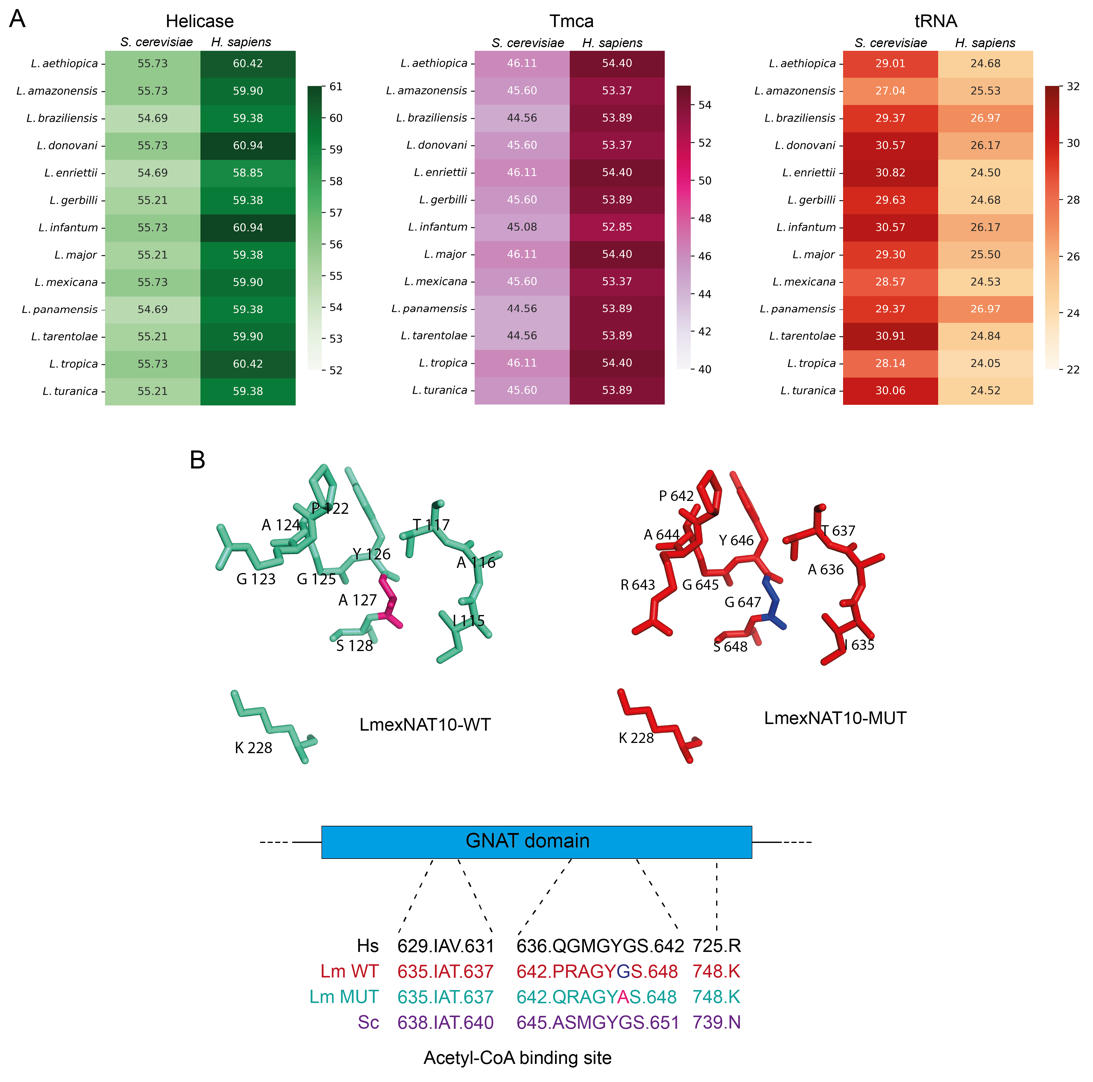

**Figure S1. Sequence Identity of NAT10 and domains among different species of *Leishmania*, *S. cerevisiae*, and humans.** **A.** The heatmaps show that the Helicase domain from *L. mexicana* and *S. cerevisiae* share 55.73% sequence identity, while *L. mexicana* and human NAT10 share ~60%. The Tmca domain of *L. mexicana* and humans, is 53.37%. Finally, the heatmap of the tRNA domain shows that there is 28.57% identity between *L. mexicana* and *S. cerevisiae*, and 24.53% when compared to the human protein domain. **B.** The amino acid change between the wild type and mutated version of NAT10 causes a subtle alteration in the structural conformation of the GNAT domain residues. The GNAT domain residues present in the Acetyl-CoA binding site are conserved among *L. mexicana* (Lm), *S. cerevisiae* (Sc) and human (Hs).

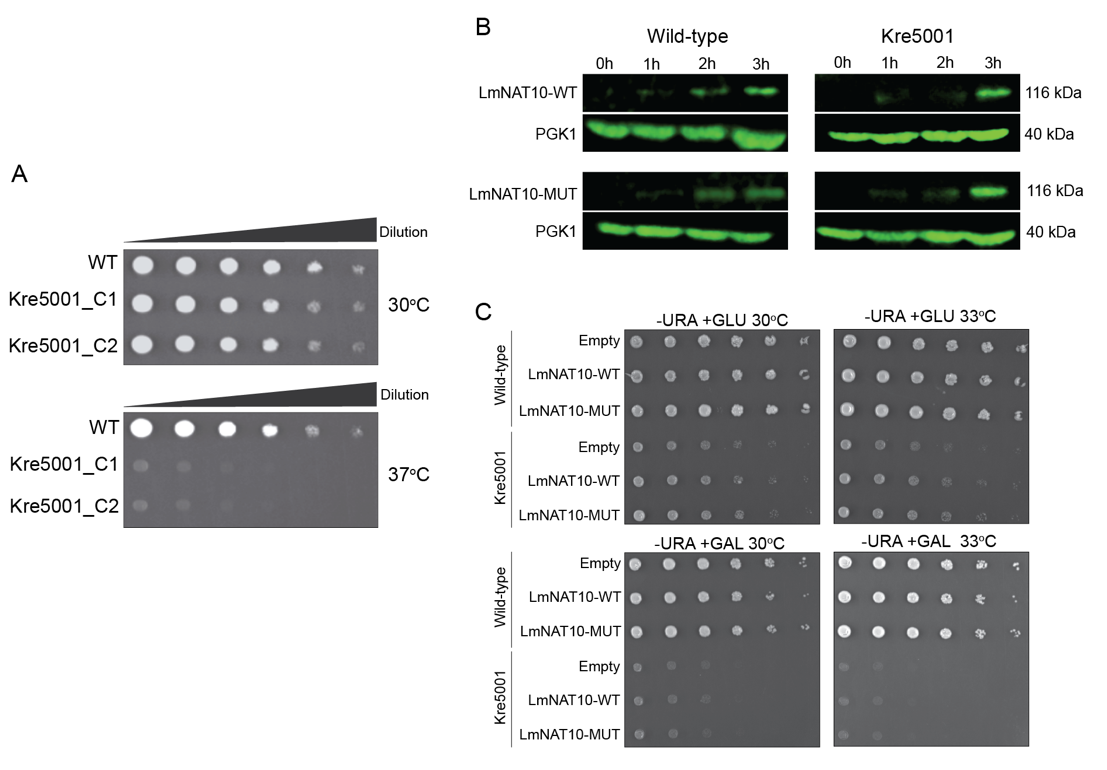

**Figure S2. *L. mexicana* NAT10 yeast complementation assays. A.** Wild type and Kre5001 *S. cerevisiae* strains were spotted on YPD solid medium and grown at 30°C and 37°C for 2-3 days. Growth of Kre5001 at 37°C was impaired compared to wild type. Similar growth rate was observed among the two cell lines at 30°C. **B.** Both *S. cerevisiae* strains (WT and Kre5001) were transfected with pYES-LmexNAT10-WT and pYES-LmexNAT10-MUT plasmids. Colonies were grown in liquid YPD medium containing raffinose and the plasmids were induced by galactose. Four-time points were collected (0, 1, 2, 3h) after galactose induction and Western Blotting was performed with an anti-FLAG tag antibody to confirm the expression of *L. mexicana* NAT10 protein. PGK1 protein was used as endogenous control. **C.** Thermosensitivity assays of the Kre5001 strain containing the plasmids pYES-LmNAT10-WT and pYES-LmNAT10-MUT. The Kre5001 strains pYES-LmNAT10-WT and pYES-LmNAT10-MUT were grown on solid medium in the presence or absence of galactose (GAL) at two temperatures 30ºC and 33ºC to assess the functional complementation of LmexNAT10. No functional complementation was observed at either temperature tested.

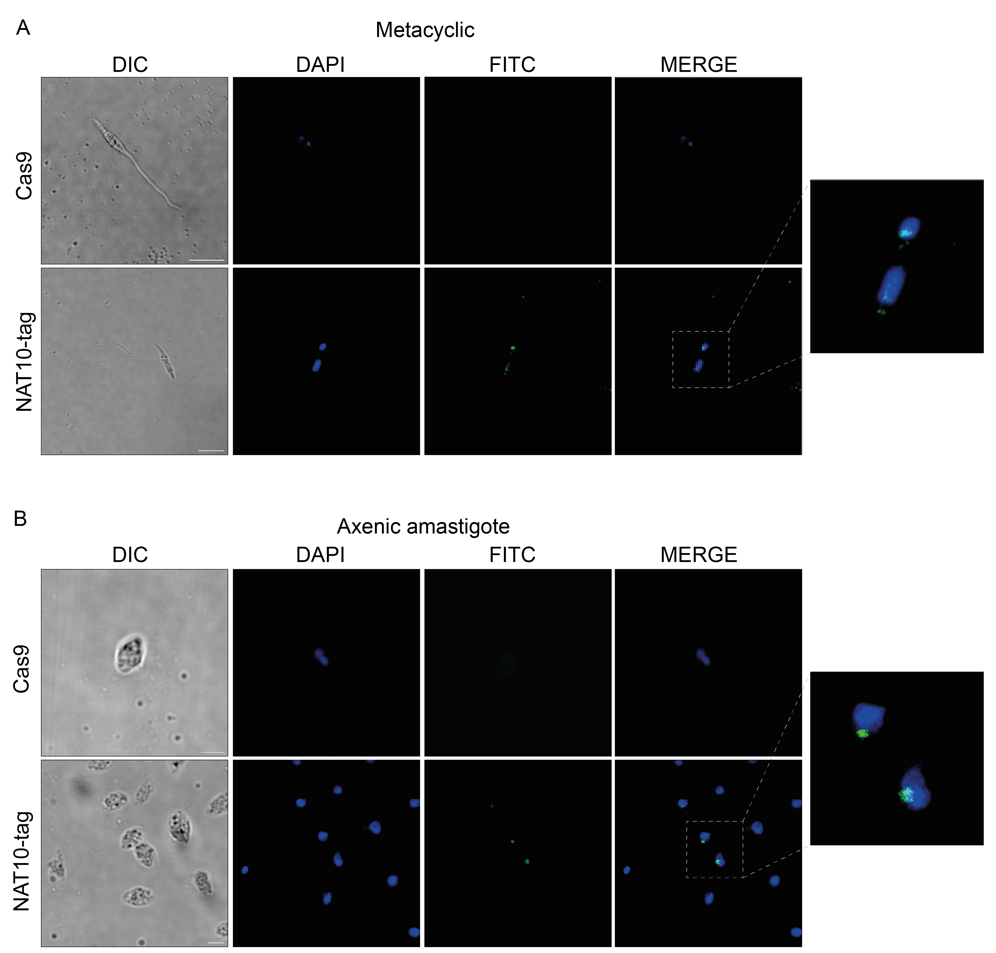

**Figure S3. *L. mexicana* NAT10 is nuclear in metacyclic and amastigote stages.** Confocal microscopy NAT10-tag cell lines of metacyclic (A) and axenic amastigote stages (B) of *L. mexicana*. Scale bars are 5 μm (metacyclic) and 2 μm (axenic amastigote).

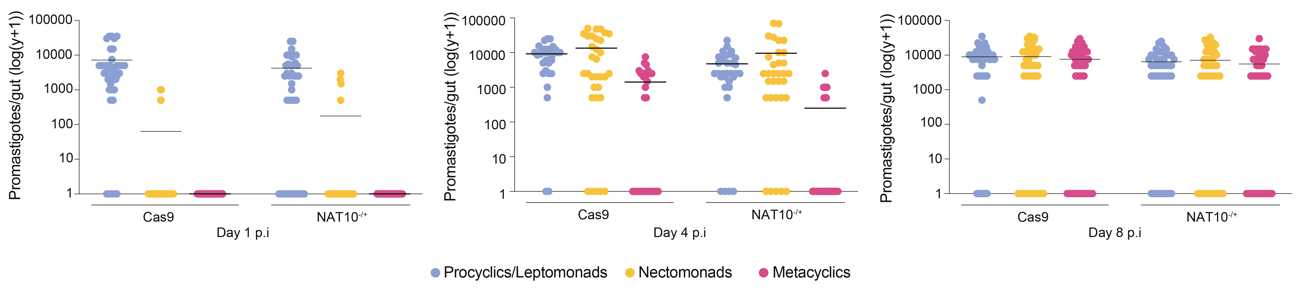

**Figure S4. *L. mexicana* NAT10^-/+^ metacyclogenesis *in vivo*.** Quantification of metacyclogenesis of Cas9 and NAT10^-/+^ at days 1, 4 and 8 post-infection of *L. longipalpis*. All infections were done at least three times with ~100 *L. longipalpis* females for each cell line.

**Supplementary Tables**

Table S1. List of primers used the homologous recombination (HR) and sgRNA fragments

| **Gene** | **Sequences** | **Primers name** |
| --- | --- | --- |
| NAT10 (LmxM.17.1250) | gaaattaatacgactcactataggGAGTGGAGGCGGGCTATACGgttttagagctagaaatagc | 5' sgRNA |
|  | gaaattaatacgactcactataggGACAGGCGCAGAATGCGATGgttttagagctagaaatagc | 3' sgRNA |
|  | ACTTGCCGGTGAACCTGCCCTTCTCTTCCCTAATACGACTCACTATAAAACTGGAAGCGCTCCTTCCTTgtataatgcagacctgctgc | Upstream forward |
|  | TGTGAAGGTACAGCGAGATACCCCCGTGGCccaatttgagagacctgtgc | Downstream reverse |
|  | GGAGCCGGCCACAGATGCCAACGACGACATactacccgatcctgatccag | Upstream reverse |

Table S2. List of PCR primers for knockout parasites confirmation

| **Gene** | **Sequences** |
| --- | --- |
| NAT10 (LmxM.17.1250) | ATCGGCGCGCGGACCGTAT  ACGTCCCTTTGTGCCTCGGTA |
| BSD resistence | GAATCCACCCTCATTGAAAGAGC  CACTATCGCTTTGATCCCAGGA |
| PAC resistence | ATGACTGAATACAAGCCAACGGTTC  CACCATGTCCTCGGCGGTTCA |

Table S3. List of RT-qPCR primers for knockout parasites confirmation

| **Gene** | **Sequences** |
| --- | --- |
| NAT10 (LmxM.17.1250) | AACGTCAAGGAGAAGCTGAAG  ACGGTTTGCATAAGGGACAG |
| GAPDH (LmxM.29.2980) | AGGAGATCGACAAGGCCATCAAGA  ACGACACGATCTTGAAGAACCGCT |
| SL-20 (Spliced leader) | ACAGTTTCTGTACTATATTG |
